# Automated DNA-Encoded Library Synthesis on Magnetic Anion Exchange Resin

**DOI:** 10.64898/2026.10.21.760614

**Authors:** Valerie Cavett, Santino L. DeGirolamo, Erika M. Cerna Arroyo, Robert Damoiseaux, Brian M. Paegel

**Affiliations:** Department of Pharmaceutical Sciences & Center for Neurotherapeutics, University of California, Irvine, California 92697, United States; Department of Chemistry, University of California, Irvine, California 92697, United States; California NanoSystems Institute and Departments of Molecular and Medical Pharmacology & Bioengineering, University of California, Los Angeles, California 90095, United States; Department of Biomedical Engineering, University of California, Irvine, California 92697, United States

## Abstract

DNA-encoded libraries (DELs) access vast chemical spaces for drug discovery, but split-and-pool DEL synthesis remains the sole and very manual portion of the DEL workflow. Here, we automate DEL construction using diethylaminoethyl-functionalized weak anion exchange (WAX) magnetic resin and a KingFisher Apex particle-handling instrument. DNA encoding occurs in solution, followed by selective immobilization of DNA conjugates under low-salt conditions that exclude > 95% of unused encoding oligonucleotides. On-DNA chemistry then proceeds using the WAX resin-adsorbed “WAX-ON” DNA substrate in neat organic solvent. The platform supports amide formation, reductive amination, copper-catalyzed azide–alkyne cycloaddition, and Fmoc removal. Automated bead transfer enables iterative reaction and washing cycles. We synthesized two-cycle 9,216-member DELs from 96 Fmoc-protected amino acids and 96 carboxylic acids. Control selections reproducibly enriched the carbonic anhydrase ligand sulfamoylbenzoic acid. Magnetic WAX-ON DEL synthesis is compatible with diverse chemistries and uses the same instrumentation to automate DEL construction and affinity screening.

## Introduction

Automated, closed-loop experimentation is increasingly of interest in drug discovery given the rise of AI-guided molecular design. Realizing these ambitions requires rapid, high-throughput, iterative mapping of small-molecule interactions with macromolecules and living cells. DNA-encoded chemical library (DEL) synthesis and screening^1,2^ together provide a key technology for realizing these ambitions for two reasons: 1) the breadth of new chemical space accessible via parallel split-and-pool synthesis,^3,4^ and 2) the density of structure-enrichment data acquired from affinity selections of libraries containing 10^6^–10^9^ different DNA-barcoded small molecules.^5,6^ These rich data sets have fueled the development of machine learning for novel building block and hit prediction.^7–9^ However, automating DEL synthesis is complicated by the many parallel chemical coupling reactions, enzymatic DNA ligations, and repeated ethanol precipitations required for DNA-encoded split-and-pool synthesis. Integrating precipitation, sample pooling, and repeated exchange between aqueous and organic reaction media complicates automated liquid handling workflows, inspiring early work on automating reagent dispensing and DNA-linked product purification.^10,11^

Reversible non-specific immobilization offers a way to simplify purification and solvent exchange by temporarily localizing DNA-linked substrates on a solid support. Halpin and Harbury demonstrated organic synthesis on reversibly immobilized, unprotected DNA oligonucleotides as part of a broader DNA-programmed synthesis platform,^12–15^ establishing a foundation for subsequent DEL chemistry development. As the field has built an expanded suite of DEL-compatible reactions, reversible immobilization via strong anion exchange resin enabled on-DNA couplings via decarboxylative cross coupling and even the highly water-sensitive stannyl amine protocol (SnAP).^14,16–18^ These pseudo-solid-phase approaches overcome DNA’s limited solubility in organic solvents, enabling reactions with highly concentrated reagents under conditions inaccessible to conventional aqueous DEL synthesis.

Solid-phase synthesis is also attractive because repetitive cycles of reaction, washing, and deprotection are readily automated. This principle underlies the automated chemical synthesis of complex biomolecules, such as peptides and oligonucleotides.^19–21^ We previously extended this concept to one-bead–one-compound DELs (OBOC-DEL),^22^ using magnetic beads and a KingFisher particle-processing instrument to automate iterative chemical synthesis and encoding.^23^ Solid-phase synthesis linker design can also be deployed to mediate the production of high-purity DELs through the ingenious dual-linker PURE-DEL approach.^24^ These studies collectively establish magnetic particle-based synthesis and handling as a viable foundation for automated DEL synthesis, employing instrumentation that already routinely automates affinity selection.^25^

Here, we combine reversible DNA immobilization with magnetic particle processing automation to establish a simplified and scalable platform for on-DNA affinity DEL synthesis. DNA-linked substrates are reversibly immobilized on diethylaminoethyl (DEAE) magnetic weak anion exchange (WAX-ON) resin and subjected to various DEL-relevant transformations, including reactions conducted under concentrated and non-aqueous conditions. WAX resin preferentially retains longer DNA constructs while rejecting shorter unincorporated encoding oligonucleotides, thereby integrating code purification into the synthesis workflow. We used this platform and a KingFisher Apex bead handling system to construct a 9,216-member DEL (96 Fmoc-protected amino acids *×* 96 carboxylic acids) and validated its performance through affinity selections and sequencing. WAX-ON DEL synthesis transforms magnetic bead handling apparatus into a fully integrated library synthesis and screening automation platform.

## Results and Discussion

### Magnetic WAX resin automates nucleic acid handling and purification

Nucleic acids are readily captured on magnetic WAX DEAE resin through interaction of the negatively charged phosphate backbone and positively charged resin-bound tertiary amines.^26,27^ We conducted T4 DNA ligase-mediated enzymatic cohesive end ligations of 5’-phosphorylated amine-modified DNA headpiece (HDNA)^1^ to a 5’-phosphorylated opening sequence (OS) containing the DEL opening primer binding site. To this HDNA–OS starting material we ligated an 11-bp 5’-phosphorylated dsDNA oligonucleotide identical in configuration to our standard DEL encoding oligonucleotides.^22^ Ligation reactions were immobilized on DEAE magnetic WAX resin under conditions of various [NaCl] and DNA products remaining in the supernatant were analyzed by PAGE (Figure 1A–B). The amount of salt required for selective elution of reversibly immobilized nucleic acid was quantitated using fluorescent dye-labeled oligonucleotides; samples were eluted in [NaCl] ranging 0.2–2 M, and elution fluorescence was determined (Figure 1C). Resin loading capacity was similarly assessed by eluting dye-labeled oligonucleotide hairpins with different lengths of base-paired regions (20, 28, and 36 bp) under optimal conditions and determining pmol released per amount of resin used (Figure 1D), which is determined spectroscopically (OD_600_).

**Figure 1.**
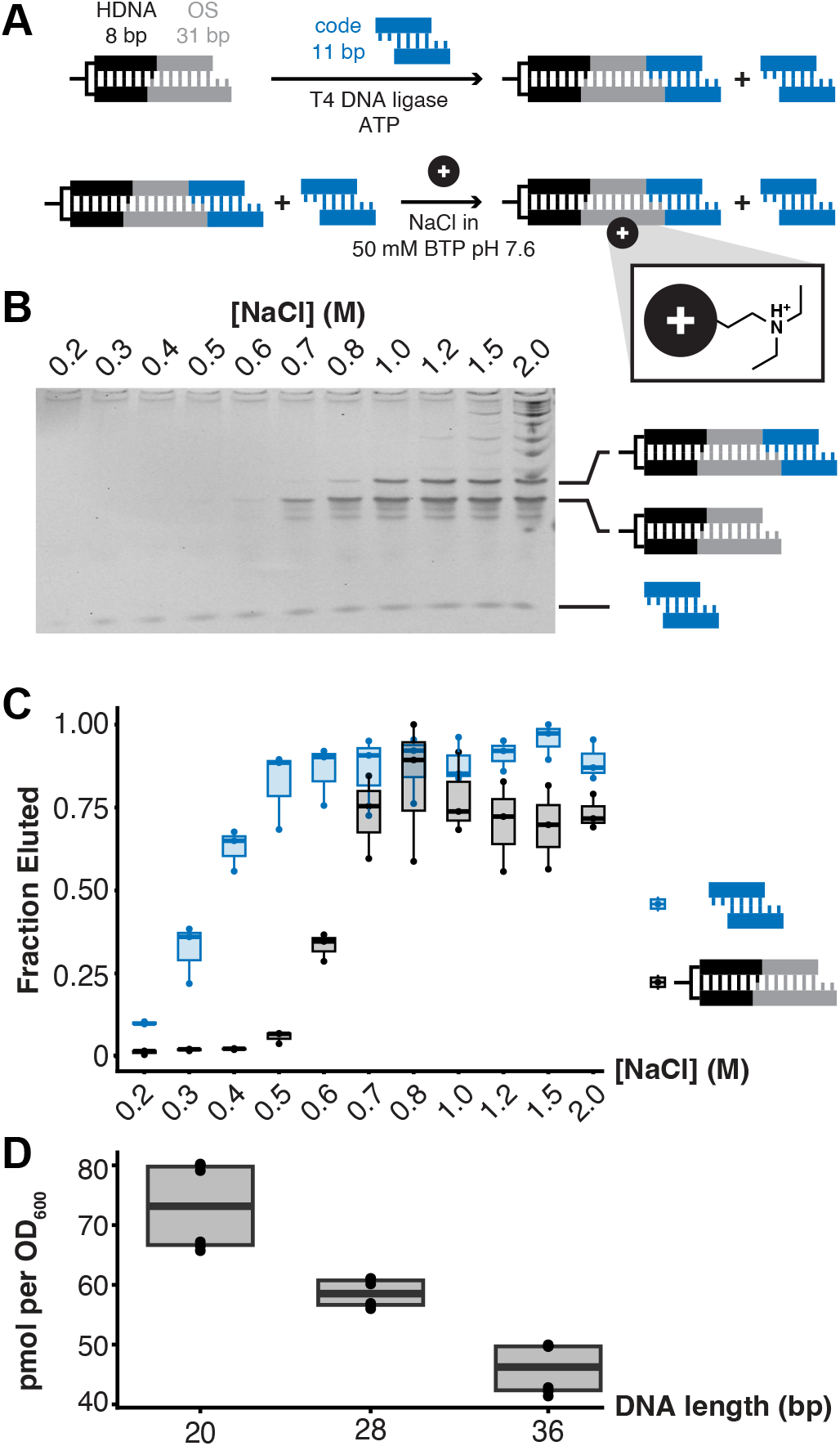
Rejection of unused coding oligonucleotides via weak anion exchange.**(A)** Synthesis begins with headpiece DNA (HDNA, black) enzymatically ligated to the opening sequence (OS, gray), followed by an 11-bp dsDNA encoding oligonucleotide (blue). **(B)** PAGE analysis of ligation reaction elutions under various [NaCl] immobilization conditions demonstrates selectivity of DEL tag binding. Addition of magnetic WAX DEAE resin in 0.3 M NaCl efficiently immobilizes the longer DEL tag, but does not bind the unincorporated 11-bp encoding oligonucleotide. Elution of fully ligated DEL tag only occurs upon addition of 1.0 M NaCl. Fluorescent dye-labeled 36-bp DNA-encoding tag (gray) or 11-bp encoding oligonucleotides (blue) were eluted from magnetic WAX DEAE resin under various [NaCl] and the supernatant was quantitated by fluorimetry. **(D)** Magnetic WAX DEAE resin loading capacity is plotted for different oligonucleotide hairpin stem lengths. Resin is quantitated via OD_600_.

Magnetic WAX DEAE resin can efficiently and selectively immobilize nucleic acids of length corresponding to the typical length of DEL encoding oligonucleotides by selecting an appropriate [NaCl]. The amount of negative charge is directly proportional to oligonucleotide length, thus separation of DEL tag from smaller nucleic acids is achieved by modulating [NaCl] and therefore shielding during resin binding. Under optimal selective binding conditions (0.5 M NaCl), the magnetic WAX resin rejects *>* 95% of unincorporated 11-bp dsDNA encoding oligonucleotides compared to the retained model 36-bp DEL tag, thus obviating expensive and manual spin column-, PAGE-, or HPLC-based sample purification steps. We hypothesized that oligonucleotide length would influence resin loading capacity. Fluorescence quantitation of eluted material of different lengths revealed a 1.5-fold drop in loading capacity per unit of resin from 22-bp to 36-bp product. The resin is far too small and irregular to count by eye or flow cytometry. An OD_600_ scattering measurement readily quantitates resin amount required for immobilization reactions.

### WAX-ON DNA chemical synthesis

On-DNA chemical synthesis was conducted using automated magnetic bead handling (KingFisher Apex, Thermo Scientific) to transfer the magnetic WAX DEAE resin through various washing, chemical reaction, and solvent exchange steps. Solid-phase DEL synthesis reaction conditions^23,28^ were employed to demonstrate various routine transformations, such as amide formation, reductive amination, Fmoc removal, and copper-catalyzed cycloaddition (CuAAC) reactions (Figure 2). All reactions were characterized via LC-MS determination of conversion yield and purity (%) based on UV absorbance (*λ* = 260 nm). Amide formation was conducted by acylation of WAX-ON HDNA primary amine with EDC-activated carboxylic acid (CA) or N-Fmoc-protected amino acid (AA) building blocks (BBs) in neat DMA solvent. Acylation reaction scope was studied using 45 CAs and 37 AAs, with 86% mean conversion yield (range 19–100%) and 82% mean purity (range 16–100%; Figures S1 and S2; Tables S5 and S6). Reductive amination was conducted on HDNA that had been WAX-ON acylated with 4-formylbenzoic acid (96% conversion yield, 100% purity, Figure S3), followed by WAX-ON imine formation and reduction with NaCNBH_3_ in MeOH (2% v/v AcOH), using the KingFisher to transfer beads sequentially from imine formation to reduction wells. Reductive amination reaction scope was studied using 12 primary amines (Am), with 96% mean conversion yield (range 87–100%) and 91% mean purity (range 85–100%; Figure S4 and Table S7). Amine deprotection and click chemistry were also explored in WAX-ON mode. Fmoc removal was quantitative using 4-methylpiperidine.^29^ CuAAC was performed by first acylating WAX-ON HDNA amine with N-Fmoc-azidoproline (quantitative conversion) followed by coupling to propargyl amine with in situ Cu(II) reduction (ascorbate) in neat DMSO with TBTA organic Cu(I) ligand.^30^ Triazole formation was quantitative with 86% purity (Figure S5).

**Figure 2.**
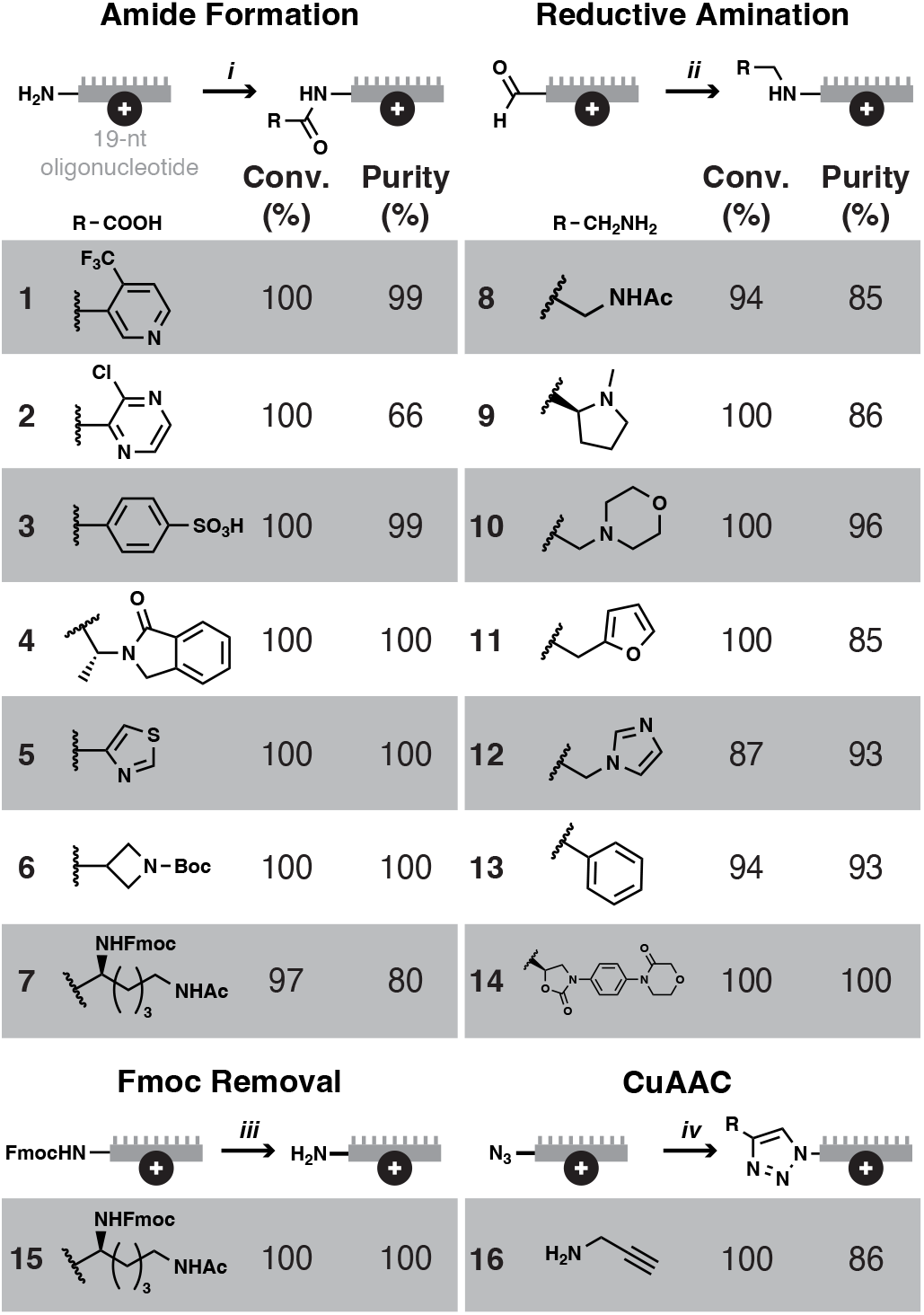
WAX-ON chemistry scope study. Reaction schemes for several DEL-relevant transformations are shown together with percent conversion yield (Conv.) and percent purity. Reaction conditions are as follows: (*i*), 50 mM AA/CA, 60 mM TMP, 5 mM HOAt, 50 mM EDC, 100 µL, 10 min, RT; (*ii*), 125 mM Am, 260 mM AcOH in MeOH, 5 min, RT; NaCNBH_3_ addition (100 µL 150 mM in MeOH), 15 min, RT; (*iii*) 4- methylpiperidine (20% v/v in DMA); (*iv*) 4 mM CuSO_4_, 20 mM ascorbate, 5 mM TBTA, 2 mM propargylamine in DMSO, 30 min, RT.

Reaction conversion yields and purity via LC-MS determination were uniformly high, with often near quantitative conversion for the transformations studied. Amide formation is a central transformation in DEL synthesis. WAX-ON chemistry enabled coupling in pure solvent, which is particularly advantageous for lipophilic BB classes such as Fmoc-protected AAs; highly lipophilic BBs can be challenging to dissolve in aqueous solvent. Reductive amination was very robust for primary amines, which benefit from acidic MeOH conditions for amine solubility as well as imine stability. Automation is further helpful for separating imine formation and borohydride-mediated imine reduction reactions. Solvent and high-yield advantages recapitulate the observations of others experimenting with solid-phase chemistry using anion exchange resins.^14,16,31^ Fmoc removal and CuAAC need not be carried out in WAX-ON mode and may not be useful in DEL synthesis, but could be very useful for completely automated post-selection on-DNA hit synthesis as on-DNA hits are popular starting points in DEL screen follow-up.^32–34^ Importantly, our exploration only encompasses some of the most popular transformations and reaction polarities; further studies of organometallic cross couplings^18,35^ and reversed reaction polarities (e.g., WAX-ON amine-DNA, reductive alkylation with aldehyde in solution) are necessary.

### Automated DEL synthesis and characterization

A two-cycle DEL was synthesized via magnetic automated WAX-ON chemistry and the resulting DEL was characterized by sequencing and affinity selection. The DEL was prepared using 96 Fmoc-protected AAs in cycle 1 and 96 CAs in cycle 2 under previously described amide formation conditions (Figure 3A).^22,36^ Enzymatic ligation of dsDNA encoding oligonucleotides was conducted in 96-well plates using the HDNA–OS starting material, then magnetic WAX resin was added and plates were transferred to the KingFisher Apex for DEL tag isolation (0.3 M NaCl), solvent exchange, and BB coupling chemistry. Cycle 1 chemistry comprised acylation with 96 N-Fmoc-protected AAs followed by Fmoc removal with 4-methylpiperidine and washing prior to pooling. The resin pool was eluted and precipitated in ethanol before the second round of encoding oligonucleotide ligation, DEL tag immobilization, and transfer to the KingFisher Apex for chemistry cycle 2 acylation with 96 CAs. The resin was then pooled for a final elution and ethanol precipitation before closing sequence (CS) ligation. Aliquots of the DEL were sequenced to assess the representation of full encoding sequences and individual BB-encoding sequences (Figure 3B). Control affinity selections were performed using biotinylated bovine carbonic anhydrase II (CAII) and structure enrichment was calculated relative to the library sequencing result (Figure 3C). Several conserved hit series in cycle 2 emerged, including **CA-33, CA-66, CA-78**, and **CA-91**.

**Figure 3.**
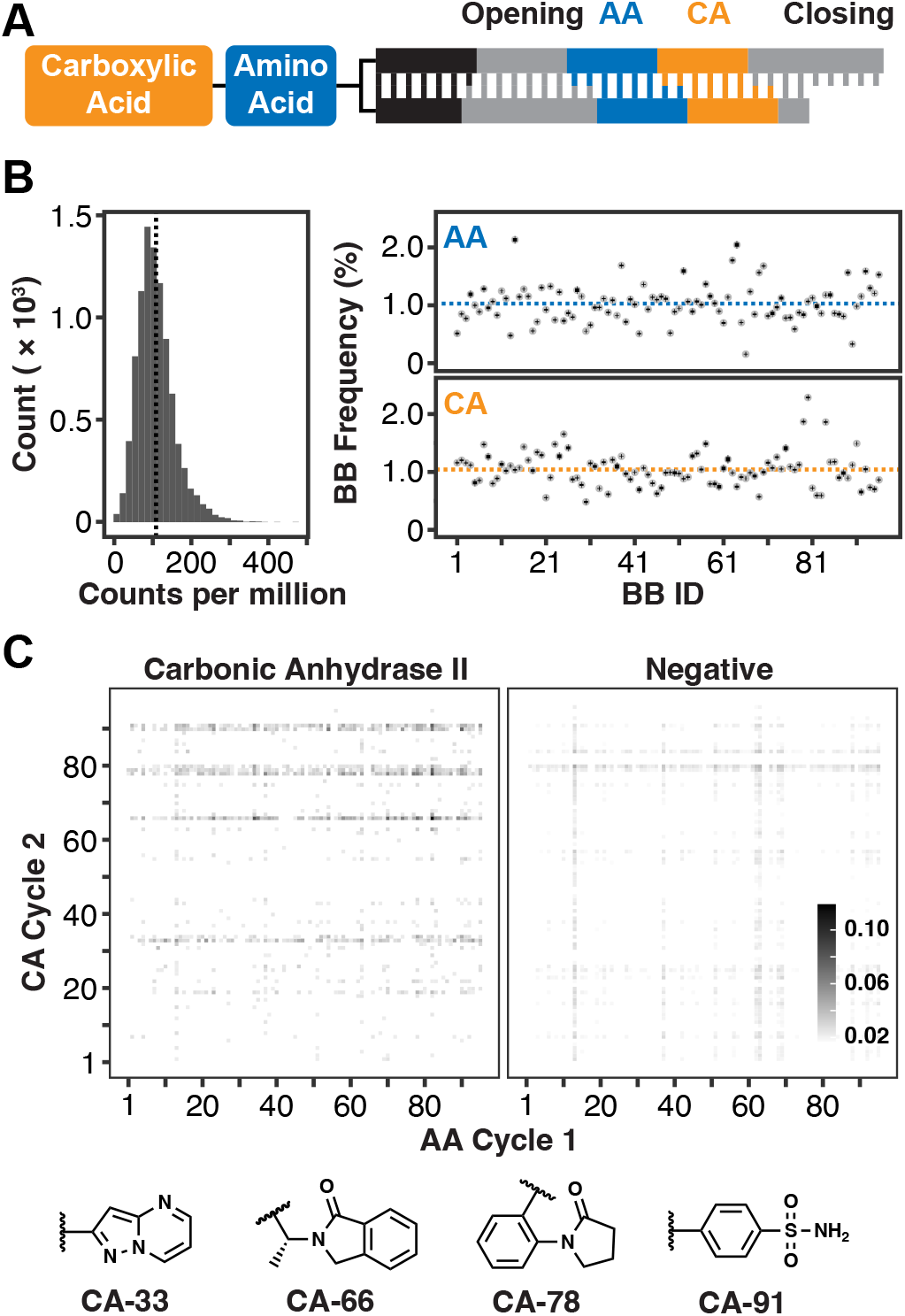
WAX-ON DEL synthesis, characterization, and implementation in affinity selection. **(A)** The DEL chemistry sequence begins in cycle 1 (blue) by acylating the HDNA–NH_2_ with 96 N-Fmoc-protected amino acids (AA). Following Fmoc removal, chemistry concludes in cycle 2 (orange) by acylating the pendant amine with 96 carboxylic acids (CA). **(B)** The unique molecular identifier (UMI) count for each library member was divided by the total UMI count in the sequenced sample to calculate the proportion of each member (vertical line indicates expected 108 per million reads mean given a 9,216-member DEL, *left*). Observed building block (BB) frequency is plotted per BB identifier (BB ID) for both cycle 1 AA and cycle 2 CA. Dashed color-coded line indicates the expected frequency based on 96 BBs in each cycle (*∼*1%). Each point is the observed BB frequency from three distinct DEL aliquots prepared for sequencing of the naive library; standard error bars are smaller than data points. **(C)** DEL member frequencies before (Negative, *right*) and after affinity selection with carbonic anhydrase II control target (CAII, *left*) were plotted as a heat map. Enriched CA structures in the CAII affinity selection are depicted.

Automated magnetic WAX-ON synthesis produced DELs with uniform code distribution and enrichment during affinity selection of ligands known to bind the established control target CAII. Automation of the complex encoded split-and-pool steps, including exchanges between different solvent systems, eliminated in-plate ethanol precipitation between ligation and chemistry steps during each synthesis cycle. Purification of reaction products or separate size exclusion steps were also eliminated. Moreover, replicate DEL syntheses are readily programmed and executed. The resulting DEL exhibits excellent code uniformity as assessed by sequencing. DEL quality control usually entails affinity selection using a protein target with well-defined binding response, such as CAII. Affinity selection using CAII as a control target enriched known ligand 4-sulfamoylbenzoic acid **91**, a cycle 2 CA, as well as pyrazolo[1,5-a]pyrimidine-2-carboxylic acid **33**. We conclude that the DEL was correctly synthesized and encoded via the automated magnetic WAX-ON protocol.

In this study, we establish magnetic WAX resin as a common substrate for enzymatic encoding, purification, and organic synthesis within an automated DEL workflow. Reversible immobilization retained full-length DEL tags while rejecting shorter encoding oligonucleotides, enabled rapid and automated exchange between aqueous and neat organic reaction media, and supported several widely used DEL transformations with high conversion. The WAX resin described here is sold as a 50 mg/mL suspension (10 mL), a mass of resin that should support a 3-cycle DEL synthesis at a 30-µmol scale according to our observed resin loading capacities, and the KingFisher quantitatively transfers 10-mg aliquots of resin (*∼*1 µmol DEL) per 96-pin magnet head, which is well within the requirements of industrial-scale library synthesis. Integration of synthesis and purification operations on a KingFisher Apex eliminated repeated in-plate precipitations that ordinarily complicate encoded split-and-pool synthesis automation and reduced each complete cycle of encoding and chemical synthesis to one day. Using this workflow, we executed syntheses of a 9,216-member DEL that exhibited uniform code distributions and reproducible enrichment of established CAII-binding chemotypes. Magnetic WAX-ON processing therefore unifies much of DEL synthesis and affinity selection on a common instrument platform while defining three important directions for further development: expanding the scope of compatible chemistry, systematically evaluating DNA compatibility, and integrating the remaining off-instrument operations into a fully automated workflow.

The chemistry demonstrated here represents a limited but useful sampling of chemical transformations adaptable to WAX-ON synthesis. Additional reaction classes, including organometallic cross-couplings, should be explored systematically. In some cases, however, the material and optical properties of the magnetic WAX resin may impose constraints on reaction performance. Photochemical and photoredox transformations, for example, could be complicated by attenuation and scattering of incident light within dense particle suspensions, particularly at the blue wavelengths commonly employed in photoredox chemistry.^37,38^ Reaction conversion alone is also insufficient to establish suitability for DEL synthesis because the conditions must preserve DNA recovery, ligation competence, and amplification fidelity.^28^ Pfizer subsequently developed a DNA-compatibility assay that couples chemical exposure with ligation and qPCR to assess the post-reaction ligation and amplification competence of the DNA tag.^33^ Incorporating such an assay into the automated WAX-ON workflow could provide a scalable means of simultaneously optimizing chemical conversion and DNA compatibility across large matrices of reaction conditions.

Automated magnetic particle handling more broadly provides the foundation for a fully integrated DEL synthesis, selection, and follow-up platform. The KingFisher Apex proved sufficient to automate nearly all of the synthesis and selection procedures described here. Because troughs are not currently available for the Apex, a KingFisher Duo Prime—the instrument commonly used to automate DEL affinity selections^25^—can alternatively perform row-by-row pooling. Looking forward, instruments designed for robotic-deck integration, such as the KingFisher Presto, could combine magnetic particle processing with automated building-block dispensing, reagent management, and plate handling. Such integration would enable automated synthesis and selection of focused affinity DELs for hit follow-up. Moreover, because our recently disclosed magnetic OBOC-DEL synthesis workflow uses an analogous particle-handling architecture,^23^ the same platform could ultimately support the synthesis of focused OBOC sub-libraries for activity-based screening.^39^

## Supporting information

DEL coding table

## Acknowledgement

This work was supported by a grant award from the National Institutes of Health (GM140890) to B.M.P. and the support of the UC Irvine Chao Family Cancer Center Genomics Research & Technology Hub (GRT Hub) for access to NGS technology (CA062203, AR075047).

## Supporting Information Available

The Supporting Information is available free of charge at the journal’s website.

- Experimental protocols
- Detailed materials manifest
- Additional results
- Building block SMILES and encoding
- KingFisher methods

## TOC Graphic

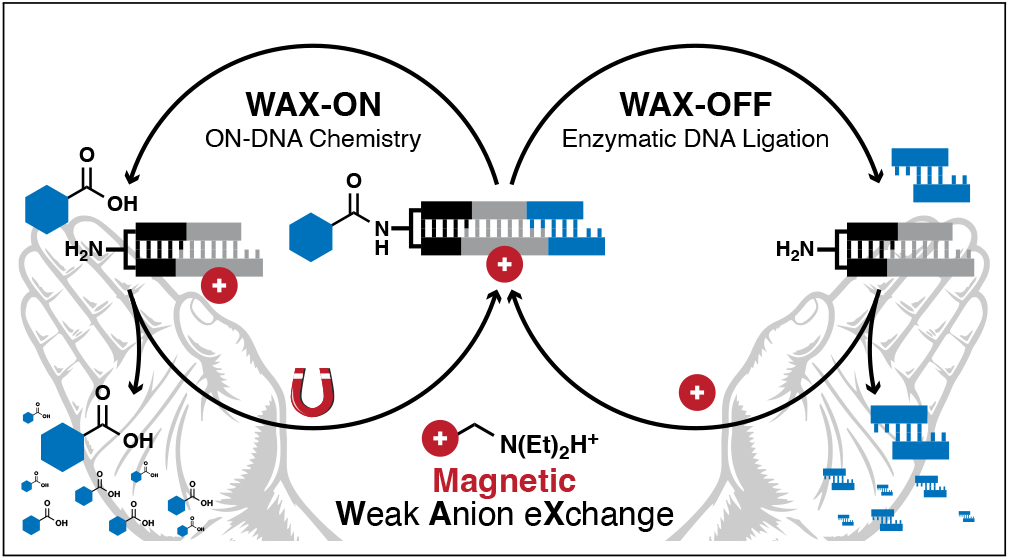

### Synopsis

Magnetic weak anion exchange resin enables automated DNA-encoded library synthesis, retaining DEL tags while rejecting unused blocks and codes.

