## Supporting Information for "Automated DNA-Encoded Library Synthesis on Magnetic Anion Exchange Resin"

#### Materials

All reagents and materials were purchased from Sigma-Aldrich (St. Louis, MO) unless otherwise specified.

- T4 DNA ligase (New England Biolabs, M0202L)
- $N,N$-Dimethylacetamide (DMA, 270555)
- 1-Hydroxy-7-azabenzotriazole (HOAt; APExBIO, A7024)
- 1-Ethyl-3-(3-dimethylaminopropyl)carbodiimide (EDC; AA Blocks, AA0033DK)
- Sodium cyanoborohydride ($NaBH_3CN$, 156159)
- Tris(benzyltriazolylmethyl)amine (TBTA, 678937)
- Copper(II) sulfate ($CuSO_4$, C1297)
- Micro Bio-Spin P-30 columns (Bio-Rad, 7326223)
- Magnetic DEAE WAX resin (1-µm; BioClone, FM102)
- HighPrep PCR Cleanup System beads (MagBio Genomics, AC-60005)
- SYBR Green I nucleic acid gel stain (Invitrogen, S7563)
- Q5 Hot Start High-Fidelity DNA polymerase (New England Biolabs, M0493L)
- Herring sperm DNA (Promega, D1815)
- Dynabeads MyOne Streptavidin C1 (Invitrogen, 65001)
- EZ-Link NHS-Biotin (Thermo Scientific, 20217)
- triethylamine (TEA, ChemImpex, 00319)
- 2,4,6-Trimethylpyridine (TMP, 142387)

- 4-formylbenzoic acid (4FBA, Ambeed, A112685)
- (2S,4R)-Fmoc-L-Pro(4-$N_3$)-OH (azidoproline, Iris Biotech GmbH, FAA3000)
- Dimethyl sulfoxide (DMSO, 472301)
- Methanol (MeOH, Fisher, A454)
- Acetic acid (AcOH, Fisher, S25118)
- 4-Methylpiperidine (4-MePip, Thermo Scientific, L02709.AE)
- Ascorbic acid (Acros Organics, 401471000)
- Hexafluoroisopropanol (HFIP, Advanced ChemTech, A0RA8406-1Z)
- Bis-Tris propane (BTP, Research Products International, B78000)
- Propargylamine (P50900)
- PLRP-S column (300 Å, 3 µm, 2.1 × 50 mm; Agilent, PL1912-1301)
- KingFisher 96-well plate (Thermo, 95040450B)
- KingFisher Duo 12 Tip Comb for 96-well plate (Thermo, 97003500)
- KingFisher 96 Tip Comb for DW Magnets (Thermo, 97002534B)
- PEG 8000 (RPI, P48080)
- Bovine carbonic anhydrase II (CAII, C2522)
- Human serum albumin (HSA, A1653)

#### Buffers

Bis-Tris propane oligonucleotide-pair buffer (BTPOP; 50 mM NaCl, 1 mM Bis-Tris propane, pH 7.6), 10 × Bis-Tris propane ligation buffer (10 × BTPLB; 500 mM NaCl, 100 mM $MgCl_2$, 10 mM ATP, 0.2% Tween-20, 100 mM Bis-Tris propane, pH 7.6), WAX immobilization buffer (WAX-ON; 20 mM sodium phosphate, pH 6.5), WAX wash buffer (WAX-WASH; 300 mM NaCl, 20 mM sodium phosphate, pH 6.5), WAX elution buffer (WAX-OFF; 1.5 M NaCl, 20 mM sodium phosphate, pH 8.3), low-salt WAX elution buffer (TEA-OFF; 10 mM TEA), PEG binding buffer (20% w/v PEG 8000, 2.5 M NaCl, 1 mM EDTA, 0.05% Tween-20, 10 mM Tris, pH 8.3), and PCR elution buffer (EB; 10 mM Tris, pH 7.5) were prepared in DI $H_2O$. Phosphate-buffered saline containing Tween-20 (PBST; 0.05% Tween-20 in PBS), biotin-supplemented PBST (PBST-B; 100 µM biotin, 0.05% Tween-20 in PBS), and herring sperm DNA-supplemented PBST (PBST+DNA; 10 µg/mL herring sperm DNA, 0.05% Tween-20 in PBS) were prepared in PBS.

#### Oligonucleotides

Oligonucleotides were purchased as desalted lyophilates and used without further purification unless otherwise specified (Integrated DNA Technologies, Inc., Coralville, IA).

Oligonucleotide nomenclature follows our previously described formalism.[S1] Briefly, all oligonucleotide sequences are written in the 5′-to-3′ direction. Oligonucleotide ligation substrates were 5′-phosphorylated (/5Phos/). Amino-modified headpiece DNA (HDNA; /5Phos/GAGTCA/iSp9//iUniAmM//iSp9/TGACTCCC) was HPLC-purified by the manufacturer and used without further purification.

Diversity-encoding oligonucleotides are designated by (≈) and four-digit numeric identifier, followed by [+] or [-] strand designation. The first digit identifies the oligonucleotide sequence set; sets 1 and 2 contain synthesis-encoding sequences. The second digit identifies the position within the encoding region, and the third and fourth digits index the individual coding sequences within each set. The common modules that install the PCR primer-binding regions flanking the synthesis encoding region are designated opening sequence (≈**OS**) and closing sequence (≈**CS**). For all modules, [+] identifies the top strand, [-] identifies the bottom strand, and [±] identifies the annealed heteroduplex formed from the corresponding [+] and [-] strands. Each annealed heteroduplex constitutes an oligonucleotide pair (OP) that serves as a substrate for enzymatic ligation. For example, annealing ≈**1101**[+] and ≈**1101**[-] yields the OP encoding module ≈**1101**[±].

OP stock solutions were prepared by combining complementary [+] and [-] strands in BTPOP (60 µM each), heating (5 min, 60 °C), and cooling to RT (5 min) before use. Encoding-module oligonucleotide sequences are provided in the Supporting Information file `DEL_encoding.csv`. All other oligonucleotide sequences are listed in Table S4.

### Methods

#### WAX Resin Characterization

**WAX resin suspension preparation and standardization.** Magnetic DEAE WAX resin (1 µm, 1 mL) was combined with WAX-WASH buffer (29 mL). An aliquot of the resin suspension was diluted 100-fold in WAX-WASH buffer, and the apparent optical density at 600 nm ($OD_{600}$) was measured (NanoDrop 2000c, Thermo Scientific). The volume of the undiluted suspension was adjusted with WAX-WASH buffer to yield a final $OD_{600}$ = 20. For volumetric measurement of resin quantities, an aliquot of resin suspension was diluted to an $OD_{600}$ of 0.1–1.0, and the dilution-adjusted $OD_{600}$ was multiplied by the suspension volume to yield the resin quantity in units of $OD_{600}\cdot$mL.

**Oligonucleotide immobilization and elution.** A mixture of a crude starting material (≈**HDNA+OS**, 100 pmol, 36 bp) and position 1 ligation product (50 pmol, 47 bp) was immobilized on resin (10 $OD_{600}$·mL). Resin was distributed to wells of 96-well plate, magnetically isolated and the supernatant removed. The immobilized oligonucleotides were eluted with NaCl solutions (15 µL; 0.1–2.0 M NaCl in 50 mM Bis-Tris propane, pH 7.6). Aliquots of each fraction were analyzed by PAGE (12%, 4 W, 25 min).

The 5′-FAM-labeled encoding module was assembled as described using ≈**11BP**[+] and ≈**11BP**[-] strands (2 nmol each in 10 µL final) to afford ≈**11BP**[±]. The ≈**11BP**[±] stock was diluted in 50 mM Bis-Tris propane (400 µL, pH 7.6). Magnetic DEAE WAX resin (420 µL; 71.6 $OD_{600}$·mL) was magnetically isolated, and the supernatant was removed. The ≈**11BP**[±] solution (400 µL, 2 nmol) was added to the resin and incubated (2 min, RT). The resin was magnetically isolated, the supernatant was removed, and the resin was washed with DI $H_2O$ (400 µL). The resin was resuspended in DI $H_2O$ (400 µL), distributed among the wells of a 96-well plate (10 µL per well), and magnetically isolated. FAM-labeled 36-bp hairpin ≈**36BP_hp** (2 nmol) was diluted in 50 mM Bis-Tris propane (800 µL, pH 7.6), heated (5 min, 95 °C), and cooled slowly to RT. Magnetic DEAE WAX resin (1 mL; 71.6 $OD_{600}$·mL) was magnetically isolated, and the supernatant was removed. The annealed hairpin solution (750 µL, 1.9 nmol) was added to the resin and incubated (2 min, RT). The resin was magnetically isolated, the supernatant was removed, and the resin was washed with DI $H_2O$ (800 µL). The resin was resuspended in DI $H_2O$ (800 µL), distributed among the wells of a 96-well plate (20 µL per well), and magnetically isolated. The immobilized oligonucleotides were eluted in triplicate with NaCl solutions (15 µL; 0.1–2.0 M NaCl in 50 mM Bis-Tris propane, pH 7.6). Eluted oligonucleotides were quantitated by fluorescence (CLARIOstar Plus, BMG LABTECH, Ortenberg, Germany; $\lambda_{ex}/\lambda_{em} = 490/530$ nm). The fraction eluted at each NaCl concentration was calculated.

**Oligonucleotide length-dependent loading capacity.** The 5′-FAM-labeled oligonucleotide hairpins, ≈**14BP_hp**, ≈**28BP_hp**, and ≈**36BP_hp** (100 pmol), were individually combined with magnetic DEAE WAX resin (1 or 0.1 $OD_{600}$·mL) in WAX-ON buffer (200 µL). Samples were mixed briefly with vortexing, incubated (3 min, RT), and the resin was magnetically isolated. The unbound fractions were collected, and the resin was washed once with WAX-WASH buffer (200 µL, 3 min, RT). Immobilized oligonucleotides were eluted with WAX-OFF buffer (200 µL, 3 min, RT). Fluorescence of the input, unbound, wash, and elution fractions was measured in triplicate ($\lambda_{ex}/\lambda_{em} = 490/530$ nm; CLARIOstar, BMG Labtech). The amount of each oligonucleotide immobilized on the resin and subsequently recovered was quantitated using a standard curve of dye-labeled oligonucleotide, yielding a length-normalized loading capacity of 1700 pmol·bp per $OD_{600}$·mL. This value is then used to determine the amount of resin required to immobilize the growing nascent DEL tag. As an example, the cycle 1 ligation product (59 bp) at 100 pmol per well (5900 pmol·bp) would require a minimum $OD_{600}$·mL of 3.5. We round this number to 4 to provide a modest excess capacity.

#### On-Resin Reaction Conversion Analysis

**General oligonucleotide immobilization and recovery procedure.** Amino-modified oligonucleotide ≈**RXN** (6 nmol) was diluted in WAX-ON (500 µL) and added to magnetically isolated DEAE WAX resin (1.3 mL of suspension, 63.9 $OD_{600}$; 83 $OD_{600}$·mL). The suspension was vortexed and incubated (3 min, RT). Aliquots of the resin suspension (100 µL; 500 pmol ≈**RXN**) were distributed into wells for automated reaction processing on a KingFisher magnetic particle processor (KingFisher Apex, Thermo Scientific). Following each reaction, the resin was washed with DMA (3 × 100 µL). For LC-MS analysis, the resin-bound oligonucleotide was eluted with TEA-OFF (100 µL). The eluates were transferred to clean microcentrifuge tubes and centrifuged (3 min, 14,000 × $g$) to pellet residual particulates. An aliquot of each clarified eluate (30 μL) was transferred to a 96-well plate for LC-MS analysis.

**Acylation.** Carboxylic acids (25 μL, 200 mM in DMSO) or Fmoc-protected amino acids (50 μL, 100 mM in DMA) were combined with TMP (25 μL, 250 mM in DMA) and HOAt (5 μL, 100 mM in DMA) in a 96-well plate. EDC (50 μL, 100 mM in DMA) was added to the monomer premix during a programmed instrument protocol pause immediately before introduction of the resin. Resin-bound ≈**RXN** was transferred to the acylation mixtures and incubated (10 min, RT). The resin was recovered, washed, and eluted according to the general procedure.

**Fmoc deprotection.** Resin-bound Fmoc-protected ≈**RXN** (500 pmol) was transferred to 20% 4-methylpiperidine in DMA (100 μL) and incubated with mixing (5 min, RT). The resin was recovered, washed, and eluted according to the general procedure.

**Reductive amination.** ≈**RXN** (20 nmol) was combined with WAX-ON buffer (1.0 mL) and magnetic DEAE WAX resin (72 $OD_{600}$·mL), and incubated with vortexing (3 min, RT). The resin was collected and the supernatant discarded. 4FBA (200 mM in DMSO, 200 μL), TMP (250 mM in DMA, 200 μL), HOAt (100 mM, 40 μL), and EDC (100 mM, 400 μL) were combined and added to the immobilized oligonucleotide. The reaction was incubated with rotation (10 min, RT, 45 rpm). The resin was collected and washed with DMA (3 × 1 mL), $H_2O$ (2 × 1 mL), and DMA (1 mL). The acylation and wash sequence was repeated twice more for a total of three acylation reactions. Resin-immobilized aldehyde-modified ≈**RXN** was aliquoted (500 pmol per reaction). Amine building block (500 mM in DMSO, 25 μL) was combined with 2% AcOH in MeOH (75 μL) and resin-immobilized aldehyde-modified ≈**RXN**. The reaction was incubated with mixing (5 min, RT). The KingFisher Apex method was paused, $NaBH_3CN$ (150 mM in MeOH, 100 μL) was added to each reaction, and the method was resumed. The resin was incubated with mixing (15 min, RT). The resin was recovered, washed, and eluted according to the general procedure.

**Copper-catalyzed azide–alkyne cycloaddition.** Amino-modified oligonucleotide ≈**RXN** was first acylated on magnetic DEAE WAX resin with Fmoc-protected azidoproline according to the general acylation procedure described above, but was not eluted from the resin. Resin-immobilized azide-modified ≈**RXN** was aliquoted (500 pmol per reaction). $CuSO_4$ (10 µL, 30 mM in DMSO), ascorbic acid (50 µL, 30 mM in DMSO), TBTA (14 µL, 30 mM in DMSO), and propargylamine (5 µL, 30 mM in DMSO) were combined sequentially and added to resin-immobilized azide-modified ≈**RXN**. The reaction was incubated with mixing (30 min, RT). The resin was recovered, washed, and eluted according to the general procedure.

**Analytical LC-MS and conversion quantitation.** Oligonucleotide reaction products were separated by reversed-phase HPLC (UltiMate 3000 UPLC, Thermo Scientific, Waltham, MA) with gradient elution (mobile phase A, 400 mM HFIP and 15 mM TEA in $H_2O$; mobile phase B, MeOH) at constant flow rate (0.35 mL/min) using a PLRP-S column (300 Å, 3 µm, 2.1 × 50 mm, Agilent) maintained at 60 °C. The standard gradient was 5% B (0–0.5 min), 5–35% B (0.5–7 min), 35–85% B (7–8.5 min), 85% B (8.5–10.5 min), 85–5% B (10.5–11 min), and 5% B (11–15 min). For analysis of products containing Fmoc-protected amino acid building blocks, an alternate gradient was used: 5% B (0–0.5 min), 5–85% B (0.5–7 min), 85–95% B (7–8.5 min), 95% B (8.5–10.5 min), 95–5% B (10.5–11 min), and 5% B (11–15 min). Eluting oligonucleotides were detected by absorbance using a diode array detector (λ = 260 nm, UltiMate 3000, Thermo Scientific) and ionized by heated electrospray. Resulting negative ions were analyzed as total ion current and spectrally resolved (150–2000 $m/z$) using a single-quadrupole mass spectrometer (ISQ EM, Thermo Scientific). Reaction conversion and purity were determined from the ratio of areas under the curve (AUC) of chromatographic features in the UV absorbance trace.

#### Automated WAX-ON DEL Synthesis

**DEL starting material preparation.** Opening sequence oligonucleotides ≈**OS**[+] and ≈**OS**[-] (0.5 mM, 20 µL each) were combined in BTPOP buffer, heated (5 min, 85 °C), and cooled to 20 °C at 0.1 °C/min to yield ≈**OS**[±]. ≈**OS**[±] was combined with HDNA (1 mM, 12 µL), 10 × BTPLB (10 µL), T4 DNA ligase (5 µL, 2000 U), and DI $H_2O$ (33 µL). The reaction was incubated (16 h, RT) and desalted using a Micro Bio-Spin P-30 column.

**Cycle 1 encoding.** Cycle 1 encoding oligonucleotides ≈**11XX**[+] and ≈**11XX**[-] were annealed as described to yield 96 cycle 1 OP encoding modules ≈**11XX**[±]. The OP encoding modules (250 pmol, 10 µL each) were arrayed in a KingFisher 96-well plate. DEL starting material (10 nmol) was combined with T4 DNA ligase (10,000 U) in BTPLB (1×, 1 mL final volume), and aliquots (10 µL, 100 pmol) were added to the arrayed OP encoding modules. The reactions were incubated (2 h, RT). Magnetic DEAE WAX resin in WAX-WASH (250 µL, $OD_{600}$ = 20) was added to each ligation reaction, and the plate transferred to the KingFisher Apex for cycle 1 on-resin chemistry.

**Cycle 1 automated chemistry.** Cycle 1 Fmoc-protected AA building blocks (25 µL, 200 mM in DMSO) and TMP (25 µL, 250 mM in DMA) were arrayed in a KingFisher 96-well plate. Immediately before initiating the KingFisher protocol, EDC and HOAt solution (50 µL; 100 mM EDC and 10 mM HOAt in DMA) was added to each well. KingFisher 96-well plates were prepared with solvent wash (3 plates, DMA, 200 µL each well), acylation reagent described above, deprotection reagent (20% 4-MePip in DMA, 100 µL each well), and solvent for resin release (1 plate, DMA, 100 µL each well). Resin-bound cycle 1-encoded DEL was processed through the acylation and Fmoc deprotection sequence described in Table S1. Resin was released in the final step into DMA (100 µL) for pooling.

**Table S1.** KingFisher Apex on-resin acylation and deprotection sequence.

| Protocol | Plate contents | V ($\mu$L) | t (min) | Mixing speed |
|---|---|---|---|---|
| Rinse | DMA | 200 | 3 | medium |
| Acylation | Acylation reagent | 100 | 10 | medium |
| Deprotection | 20% 4-MePip in DMA | 100 | 10 | medium |
| Wash | DMA | 200 | 3 | medium |
| Wash | DMA | 200 | 3 | medium |
| Release beads | DMA | 100 | 3.5 | fast |

**Cycle 1 pooling and DEL recovery.** The resin suspensions were pooled in a clean 50-mL tube. The resin was washed with DI $H_2O$ (2 × 2 mL) followed by WAX-WASH (2 × 2 mL), and the cycle 1 DEL was eluted with WAX-OFF (2.5 mL). The eluted DEL was combined with NaOAc (250 µL, 3 M, pH 5), EtOH (12.5 mL), and linear polyacrylamide (10 µg) and incubated (2 h, -80 °C). The precipitated DEL was collected by centrifugation (20 min, 4 °C, 7,000 × $g$) and washed with 80% ice-cold EtOH (2 × 500 µL). The pellet was resuspended in DI $H_2O$ (250 µL), and the recovered DEL was quantitated by absorbance (λ = 260 nm).

**Cycle 2 encoding.** Cycle 2 encoding oligonucleotides ≈**22XX**[+] and ≈**22XX**[-] were annealed as described to yield 96 cycle 2 OP encoding modules ≈**22XX**[±]. The OP encoding modules (250 pmol, 10 µL each) were arrayed in a KingFisher 96-well plate. Cycle 1 DEL (10 nmol) was combined with T4 DNA ligase (10,000 U) in BTPLB (1×, 1 mL final volume), and aliquots (10 µL, 100 pmol) were added to the arrayed OP encoding modules. The reactions were incubated (2 h, RT). Magnetic DEAE WAX resin (250 µL, $OD_{600}$ = 20) in WAX-WASH was added to each ligation reaction, and the plate transferred to the KingFisher Apex for cycle 2 on-resin chemistry.

**Cycle 2 automated chemistry.** Cycle 2 carboxylic acid building blocks (25 µL, 200 mM in DMSO) and TMP (25 µL, 250 mM in DMA) were arrayed in a KingFisher 96-well plate. Immediately before initiating the KingFisher protocol, EDC and HOAt solution (50 µL; 100

mM EDC and 10 mM HOAt in DMA) was added to each well. KingFisher 96-well plates were prepared with solvent wash (3 plates, DMA, 200 µL each well), acylation reagent described above, and solvent for resin release (1 plate, DMA, 100 µL each well). Resin-bound cycle 2-encoded DEL was processed through the acylation sequence described in Table S2. Resin was released in the final step into DMA (100 µL) for pooling.

**Table S2.** KingFisher Apex on-resin acylation sequence.

| Protocol | Plate contents | V ($\mu$L) | t (min) | Mixing speed |
|---|---|---|---|---|
| Rinse | DMA | 200 | 3 | medium |
| Acylation | Acylation reagent | 100 | 10 | medium |
| Wash | DMA | 200 | 3 | medium |
| Wash | DMA | 200 | 3 | medium |
| Release beads | DMA | 100 | 3.5 | fast |

**Cycle 2 pooling and DEL recovery.** The resin suspensions were pooled in a clean 50-mL tube. The resin was washed with DI $H_2O$ (2 $\times$ 2 mL) followed by WAX-WASH (2 $\times$ 2 mL), and the cycle 2 DEL was eluted with WAX-OFF (2.5 mL). The eluted DEL was combined with NaOAc (250 µL, 3 M, pH 5), EtOH (12.5 mL), and linear polyacrylamide (10 µg) and incubated (2 h, -80 °C). The precipitated DEL was collected by centrifugation (20 min, 4 °C, 7,000 $\times$ $g$) and washed with 80% ice-cold EtOH (2 $\times$ 500 µL). The pellet was resuspended in DI $H_2O$ (100 µL), and the recovered DEL was quantitated by absorbance (λ = 260 nm).

**Closing sequence ligation.** Closing sequence oligonucleotides $\approx$**CS**[+] and $\approx$**CS**[-] (1 mM, 2 µL each) were annealed as described to yield the closing sequence module $\approx$**CS**[$\pm$]. $\approx$**CS**[$\pm$], 10 $\times$ BTPLB (1$\times$ final concentration), and T4 DNA ligase (2,400 U) were added to the recovered cycle 2 DEL. The reaction was incubated (16 h, RT) and desalted using a Micro Bio-Spin P-30 column.

**DEL quantitation by qPCR.** The qPCR matrix contained Q5 Hot Start High-Fidelity DNA polymerase, oligonucleotide primers $\approx$**P1** and $\approx$**P2** (0.5 µM each), dNTPs (0.2 mM),

SYBR Green I (0.2×), and Q5 reaction buffer (1×). Serial dilutions of template standard ≈**DEL_TEM** (100 pM–1 fM) were prepared by 10-fold dilution. The completed DEL was diluted 1:1,000, 1:10,000, and 1:100,000 in DI $H_2O$. Template standards (1 µL, duplicate) or DEL dilutions (1 µL, triplicate) were added to qPCR matrix (20 µL). Reactions were thermally cycled (98 °C, 1 min; [98 °C, 10 s; 60 °C, 12 s; 72 °C, 15 s] × 30 cycles) with fluorescence monitoring (QuantStudio 3, Thermo Fisher Scientific). DEL concentration was determined from the ≈**DEL_TEM** template standard curve.

#### DEL Characterization and Affinity Selection

**Target preparation.** Bovine carbonic anhydrase II (CAII) and human serum albumin (HSA) were chemically biotinylated with NHS-biotin according to the manufacturer's protocol.
**DEL affinity selection.** Biotinylated CAII or HSA (5 µM, 100 µL in PBS) was immobilized on Dynabeads MyOne Streptavidin C1 (10 µL) using a KingFisher Duo Prime (30 min, medium mixing). Target-loaded beads were washed sequentially with PBST-B and PBST as described in Table S3. DEL (0.1 pmol, 100 µL in PBST+DNA) was incubated with the target-loaded beads (60 min, medium mixing), followed by four washes with PBST. Bound DEL was released into EB (100 µL, 3 min, medium mixing).

**Table S3.** KingFisher Duo Prime DEL affinity selection protocol.

| Protocol | Plate contents | V ($\mu$L) | t (min) | Mixing speed |
|---|---|---|---|---|
| Load protein | Target | 100 | 30 | medium |
| Wash | PBST-B | 200 | 3 | medium |
| Wash | PBST | 200 | 3 | medium |
| Wash | PBST | 200 | 3 | medium |
| Selection | DEL | 100 | 60 | medium |
| Wash | PBST | 200 | 1 | medium |
| Wash | PBST | 200 | 1 | medium |
| Wash | PBST | 200 | 1 | medium |
| Wash | PBST | 200 | 1 | medium |
| Elution | EB | 100 | 3 | medium |

**NGS library preparation.** Replicate aliquots of each selected DEL population (2% of recovered material, $n = 3$) were amplified by PCR using oligonucleotide primers ≈**P1** and ≈**P2**. Illumina P5 and P7 adapters and sample-specific i5 and i7 barcodes were subsequently installed by PCR using custom adapter oligonucleotides (0.1 µM each). The PCR matrix contained Q5 Hot Start High-Fidelity DNA polymerase (0.01 U/$\mu$L), dNTPs (0.2 mM), SYBR Green I (0.2×), and Q5 reaction buffer (1×). Reactions were thermally cycled (98 °C, 1 min; [98 °C, 10 s; 75 °C, 15 s] × 7 cycles). Adapter-modified DNA was purified using HighPrep PCR Cleanup System beads and PEG binding buffer (0.7× sample volume). Beads were washed with 80% EtOH (100 µL) and purified DNA was eluted in 10 mM Tris (40 µL, pH 8.3). Adapter-modified DNA was quantitated by qPCR (KAPA Biosystems, Wilmington, MA), pooled, denatured, and diluted to 12 pM. The pooled library was sequenced using a MiSeq i100 flow cell (5 million reads, 300-cycle chemistry, dual indexing; Illumina, San Diego, CA).

**Sequence processing and enrichment analysis.** Sequencing reads were aligned to the library consensus sequence ≈**MOTIF**, requiring exact matches at each encoding region. Reads were assigned to the corresponding cycle 1 and cycle 2 building-block identifiers, and counts were aggregated for each library member.

#### Supplemental Tables

**Table S4.** Oligonucleotide sequences.

| Identifier | Description | Sequence (5′–3′) |
|---|---|---|
| **MOTIF** | Library consensus sequence | TGACTCCCGCCGCCGCCTTCGTCCTTCTCAGCGACATGGNNNNNNNNTCANNNNNNNNNGTTCCCAAACNNNNNNNNNGTTTGCCCGCCAGTTGTTGTGCCAC |
| **HDNA** | Amino-modified headpiece DNA | /5Phos/GAGTCA/iSp9//iUniAmM//iSp9/TGACTCCC |
| **OS**[+] | Opening sequence top strand | /5Phos/GCCGCCGCCTTCGTCCTTCTCAGCGAC |
| **OS**[-] | Opening sequence bottom strand | /5Phos/CCATGTCGCTGAGAAGGACGAAGGCGGCGGCGG |
| **CS**[+] | Closing sequence top strand | /5Phos/GTTCCCAAACNNNNNNNNNGTTTGCCCGCCAGTTGTTGTGCCAC |
| **CS**[-] | Closing sequence bottom strand | /5Phos/GTTTGGG |
| **P1** | qPCR forward primer | GCCGCCGCCTTCGTCCTTCTCAGCGAC |
| **P2** | qPCR reverse primer | GTGGCACAACAACTGGCGGGCAAAC |
| **P5** | Secondary PCR forward primer | AATGATACGGCGACCACCGA |
| **P7** | Secondary PCR reverse primer | CAAGCAGAAGACGGCATACGA |
| **11BP[+]** | 11-bp resin loading probe top strand | /5FAM/AGACCGAGATAG |
| **11BP[-]** | 11-bp resin loading probe bottom strand | CTATCTCGGTC |
| **20BP_hp** | 44-nt, 20-bp hairpin resin loading probe | /5FAM/CATGCACCATGGCATGCATGGTAACATGCATGCCATGGTGCATG |
| **28BP_hp** | 60-nt, 28-bp hairpin resin loading probe | /5FAM/CATGCACCATGGCATGCATGCATGCATGGTAACATGCATGCATGCATGCCATGGTGCATG |
| **36BP_hp** | 76-nt, 36-bp hairpin resin loading probe | /5FAM/CATGCACCATGGCATGCATGCATGCATGCATGCATGGTAACATGCATGCATGCATGCATGCATGCCATGGTGCATG |
| **RXN** | Reaction conversion probe | /5AmMC6/CACTCAACCCTATCTCGGT |
| **DEL_TEM** | qPCR DEL standard | GCCGCCGCCTTCGTCCTTCTCAGCGACATGGTGGAAAGTTCAAACCCTACGTTCCCAAACATCGATCCGTTTGCCCGCCAGTTGTTGTGCCAC |

**Table S5.** Acylation of immobilized nucleic acid. 500 pmol of a model $NH_2$-modified nucleic acid was immobilized on 1-µm magnetic DEAE WAX resin and acylated with a carboxylic acid (CA). After elution, samples were analyzed by HPLC monitoring UV absorbance (λ= 260 nm). Conversion was calculated as the percentage of the expected product peak area to the sum of the starting material and expected product peak areas. Purity of the product was calculated as a percentage of product peak areas among all peaks excluding starting material.

| SMILES | Monomer Class | Reaction | Conversion (%) | Purity (%) | Theoretical MW | Expected m/z | Calculated MW |
|---|---|---|---|---|---|---|---|
| O=C(O)C1=CN=CC=C1C(F)(F)F | CA | acylation | 100 | 99 | 6036.0 | z=-4: 1508, z=-5: 1206.2, z=-6: 1005, z=-7: 861.3, z=-8: 753.5 | 6036.1 |
| O=C(C1=NC=CN=C1Cl)O | CA | acylation | 100 | 66 | 6003.4 | z=-4: 1499.9, z=-5: 1199.7, z=-6: 999.6, z=-7: 856.6, z=-8: 749.4 | 6004.4 |
| O=C(C1=CN=C(N(C)N=C2C)C2=C1Cl)O | CA | acylation | 98 | 100 | 6070.5 | z=-4: 1516.6, z=-5: 1213.1, z=-6: 1010.8, z=-7: 866.2, z=-8: 757.8 | 6071.2 |
| ClC1=CC=CC(C(O)=O)=N1 | CA | acylation | 100 | 100 | 6002.4 | z=-4: 1499.6, z=-5: 1199.5, z=-6: 999.4, z=-7: 856.5, z=-8: 749.3 | 6002.6 |
| O=C(O)C1=CC=C(S(=O)([O-])=O)C=C1 | CA | acylation | 100 | 97 | 6046.1 | z=-4: 1510.5, z=-5: 1208.2, z=-6: 1006.7, z=-7: 862.7, z=-8: 754.8 | 6046.6 |
| OC(c(cccc1)c1SC(F)(F)F)=O | CA | acylation | 100 | 72 | 6067.1 | z=-4: 1515.8, z=-5: 1212.4, z=-6: 1010.2, z=-7: 865.7, z=-8: 757.4 | 6067.0 |
| OC(c(cc1)c(C(F)(F)F)cc1Cl)=O | CA | acylation | 97 | 97 | 6069.4 | z=-4: 1516.4, z=-5: 1212.9, z=-6: 1010.6, z=-7: 866.1, z=-8: 757.7 | 6068.0 |
| Cc(ccc(F)c1)c1C(O)=O | CA | acylation | 95 | 100 | 5999.0 | z=-4: 1498.8, z=-5: 1198.8, z=-6: 998.8, z=-7: 856, z=-8: 748.9 | 5999.9 |
| C[C@H](C(O)=O)N(Cc1c2cccc1)C2=O | CA | acylation | 100 | 100 | 6050.1 | z=-4: 1511.5, z=-5: 1209, z=-6: 1007.3, z=-7: 863.3, z=-8: 755.3 | 6050.6 |
| O=C(C1=CSC=N1)O | CA | acylation | 100 | 100 | 5974.0 | z=-4: 1492.5, z=-5: 1193.8, z=-6: 994.7, z=-7: 852.4, z=-8: 745.8 | 5974.1 |
| O=C(O)C1=C(C(F)(F)F)OC(C)=C1 | CA | acylation | 100 | 98 | 6039.0 | z=-4: 1508.7, z=-5: 1206.8, z=-6: 1005.5, z=-7: 861.7, z=-8: 753.9 | 6038.9 |
| O=C(C1CN(C(OC(C)(C)C)=O)C1)O | CA | acylation | 100 | 100 | 6046.1 | z=-4: 1510.5, z=-5: 1208.2, z=-6: 1006.7, z=-7: 862.7, z=-8: 754.8 | 6047.1 |
| O=C(O)c1cc(C2CC2)n[nH]1 | CA | acylation | 74 | 98 | 5997.0 | z=-4: 1498.3, z=-5: 1198.4, z=-6: 998.5, z=-7: 855.7, z=-8: 748.6 | 5997.9 |
| O=C(O)CCn1c(=O)sc2ccccc21 | CA | acylation | 95 | 85 | 6068.1 | z=-4: 1516, z=-5: 1212.6, z=-6: 1010.4, z=-7: 865.9, z=-8: 757.5 | 6069.2 |
| CN(CC(=O)O)c1nc2ccccc2s1 | CA | acylation | 91 | 78 | 6067.2 | z=-4: 1515.8, z=-5: 1212.4, z=-6: 1010.2, z=-7: 865.7, z=-8: 757.4 | 6066.7 |
| O=C(O)c1cc(C2CC2)on1 | CA | acylation | 100 | 100 | 5998.0 | z=-4: 1498.5, z=-5: 1198.6, z=-6: 998.7, z=-7: 855.9, z=-8: 748.8 | 5998.2 |
| COc1cccc(C(=O)O)c1OC | CA | acylation | 85 | 89 | 6027.1 | z=-4: 1505.8, z=-5: 1204.4, z=-6: 1003.5, z=-7: 860, z=-8: 752.4 | 6027.7 |
| Cc1nc(C(=O)O)co1 | CA | acylation | 100 | 82 | 5972.0 | z=-4: 1492, z=-5: 1193.4, z=-6: 994.3, z=-7: 852.1, z=-8: 745.5 | 5971.0 |
| CCn1nc2c(c1C(=O)O)CCC2 | CA | acylation | 83 | 74 | 6025.1 | z=-4: 1505.3, z=-5: 1204, z=-6: 1003.2, z=-7: 859.7, z=-8: 752.1 | 6025.0 |
| O=C(O)CCc1ccon1 | CA | acylation | 92 | 86 | 5986.0 | z=-4: 1495.5, z=-5: 1196.2, z=-6: 996.7, z=-7: 854.1, z=-8: 747.3 | 5986.3 |
| COc1cccc(OCC(=O)O)c1 | CA | acylation | 92 | 81 | 6027.1 | z=-4: 1505.8, z=-5: 1204.4, z=-6: 1003.5, z=-7: 860, z=-8: 752.4 | 6027.1 |
| COc1ccccc1OCC(=O)O | CA | acylation | 97 | 92 | 6027.1 | z=-4: 1505.8, z=-5: 1204.4, z=-6: 1003.5, z=-7: 860, z=-8: 752.4 | 6026.0 |
| O=C(O)c1cc2cccnc2[nH]1 | CA | acylation | 88 | 93 | 6007.0 | z=-4: 1500.8, z=-5: 1200.4, z=-6: 1000.2, z=-7: 857.1, z=-8: 749.9 | 6007.5 |
| O=C(O)c1csc2cnccc12 | CA | acylation | 80 | 98 | 6024.1 | z=-4: 1505, z=-5: 1203.8, z=-6: 1003, z=-7: 859.6, z=-8: 752 | 6024.2 |
| O=C(O)CSc1ccccc1 | CA | acylation | 81 | 87 | 6013.1 | z=-4: 1502.3, z=-5: 1201.6, z=-6: 1001.2, z=-7: 858, z=-8: 750.6 | 6012.3 |
| O=C(O)c1ccc(C(F)F)s1 | CA | acylation | 95 | 89 | 6023.0 | z=-4: 1504.8, z=-5: 1203.6, z=-6: 1002.8, z=-7: 859.4, z=-8: 751.9 | 6022.2 |
| O=C(O)c1ccc2c(n1)NCCC2 | CA | acylation | 54 | 48 | 6023.1 | z=-4: 1504.8, z=-5: 1203.6, z=-6: 1002.8, z=-7: 859.4, z=-8: 751.9 | 6023.1 |
| O=C(O)Cn1cnc(-c2ccccc2)cc1=O | CA | acylation | 94 | 28 | 6075.1 | z=-4: 1517.8, z=-5: 1214, z=-6: 1011.5, z=-7: 866.9, z=-8: 758.4 | 6075.1 |
| Cc1csc2ncc(C(=O)O)c(=O)n12 | CA | acylation | 79 | 84 | 6055.1 | z=-4: 1512.8, z=-5: 1210, z=-6: 1008.2, z=-7: 864, z=-8: 755.9 | 6054.6 |
| O=C(O)C1(c2cccs2)CCCC1 | CA | acylation | 93 | 88 | 6041.2 | z=-4: 1509.3, z=-5: 1207.2, z=-6: 1005.9, z=-7: 862, z=-8: 754.1 | 6041.1 |
| Cc1nn2cnnc2c(C)c1CCC(=O)O | CA | acylation | 98 | 95 | 6065.1 | z=-4: 1515.3, z=-5: 1212, z=-6: 1009.9, z=-7: 865.4, z=-8: 757.1 | 6065.5 |
| O=C(O)C1(n2cccc2)CCCC1 | CA | acylation | 94 | 87 | 6024.1 | z=-4: 1505, z=-5: 1203.8, z=-6: 1003, z=-7: 859.6, z=-8: 752 | 6023.8 |
| CC(C)c1cc(=O)[nH]c(SCC(=O)O)n1 | CA | acylation | 92 | 92 | 6073.2 | z=-4: 1517.3, z=-5: 1213.6, z=-6: 1011.2, z=-7: 866.6, z=-8: 758.1 | 6072.6 |
| O=C(O)c1cc2c([nH]c1=O)CCCCC2 | CA | acylation | 26 | 60 | 6052.1 | z=-4: 1512, z=-5: 1209.4, z=-6: 1007.7, z=-7: 863.6, z=-8: 755.5 | 6052.8 |
| O=C(O)C1(c2ccccc2)CCCC1 | CA | acylation | 71 | 83 | 6035.1 | z=-4: 1507.8, z=-5: 1206, z=-6: 1004.9, z=-7: 861.2, z=-8: 753.4 | 6035.4 |
| O=C(O)Cn1cnc2ccccc2c1=O | CA | acylation | 91 | 83 | 6049.1 | z=-4: 1511.3, z=-5: 1208.8, z=-6: 1007.2, z=-7: 863.2, z=-8: 755.1 | 6048.0 |
| OC(c(cc1)cc(NC2)c1NC2=O)=O | CA | acylation | 100 | 83 | 6037.1 | z=-4: 1508.3, z=-5: 1206.4, z=-6: 1005.2, z=-7: 861.4, z=-8: 753.6 | 6034.1 |
| O=C(O)c1ccc(N2CCOCC2)nn1 | CA | acylation | 85 | 91 | 6054.1 | z=-4: 1512.5, z=-5: 1209.8, z=-6: 1008, z=-7: 863.9, z=-8: 755.8 | 6053.7 |
| Cc1nn(-c2ccccc2)c(C)c1C(=O)O | CA | acylation | 19 | 88 | 6061.1 | z=-4: 1514.3, z=-5: 1211.2, z=-6: 1009.2, z=-7: 864.9, z=-8: 756.6 | 6061.6 |
| O=C(O)c1noc2c1CCCC2 | CA | acylation | 82 | 99 | 6012.1 | z=-4: 1502, z=-5: 1201.4, z=-6: 1001, z=-7: 857.9, z=-8: 750.5 | 6013.6 |
| O=C(O)c1ccc2cc[nH]c2c1 | CA | acylation | 72 | 92 | 6006.0 | z=-4: 1500.5, z=-5: 1200.2, z=-6: 1000, z=-7: 857, z=-8: 749.8 | 6006.6 |
| O=C(O)c1ccc(Oc2nccs2)cc1 | CA | acylation | 95 | 90 | 6066.1 | z=-4: 1515.5, z=-5: 1212.2, z=-6: 1010, z=-7: 865.6, z=-8: 757.3 | 6066.6 |
| COc1ccccc1C1(C(=O)O)CC1 | CA | acylation | 78 | 84 | 6037.1 | z=-4: 1508.3, z=-5: 1206.4, z=-6: 1005.2, z=-7: 861.4, z=-8: 753.6 | 6038.3 |
| O=C(O)c1ccncn1 | CA | acylation | 95 | 98 | 5969.0 | z=-4: 1491.2, z=-5: 1192.8, z=-6: 993.8, z=-7: 851.7, z=-8: 745.1 | 5970.1 |
| Cc1nc2nc(C(=O)O)cc(C(C)(C)C)n2n1 | CA | acylation | 88 | 78 | 6079.1 | z=-4: 1518.8, z=-5: 1214.8, z=-6: 1012.2, z=-7: 867.4, z=-8: 758.9 | 6078.8 |

**Table S6.** Acylation of immobilized nucleic acid. 500 pmol of a model $NH_2$-modified nucleic acid was immobilized on 1-µm magnetic DEAE WAX resin and subjected to acylation with an Fmoc-amino acid (AA). After elution from the resin, samples were analyzed by HPLC (260 nm). Conversion was calculated by comparison of the expected product peak area to the starting material. Purity of the product was calculated as a percentage of all peaks excluding any unconverted starting material.

| SMILES | Monomer Class | Reaction | Conversion (%) | Purity (%) | Theoretical MW | Expected m/z | Calculated MW |
|---|---|---|---|---|---|---|---|
| C1[C@@H](CN([C@@H]1C(=O)O)C(=O)OCC2C3=CC=CC=C3C4=CC=CC=C24)N=[N+]=[N-] | AA | acylation | 84 | 93 | 6223.3 | z=-4: 1554.8, z=-5: 1243.7, z=-6: 1036.2, z=-7: 888, z=-8: 776.9 | 6223.8 |
| C1[C@H](CN([C@@H]1C(=O)O)C(=O)OCC2C3=CC=CC=C3C4=CC=CC=C24)N=[N+]=[N-] | AA | acylation | 97 | 88 | 6223.3 | z=-4: 1554.8, z=-5: 1243.7, z=-6: 1036.2, z=-7: 888, z=-8: 776.9 | 6223.8 |
| CC(=O)NCCCC[C@@H](C(=O)O)NC(=O)OCC1C2=CC=CC=C2C3=CC=CC=C13 | AA | acylation | 97 | 80 | 6255.4 | z=-4: 1562.8, z=-5: 1250.1, z=-6: 1041.6, z=-7: 892.6, z=-8: 780.9 | 6255.1 |
| CC(C)(C)OC(=O)N1CCC(CC1)N(CC(=O)O)C(=O)OCC2C3=CC=CC=C3C4=CC=CC=C24 | AA | acylation | 69 | 90 | 6325.1 | z=-4: 1580.3, z=-5: 1264, z=-6: 1053.2, z=-7: 902.6, z=-8: 789.6 | 6325.9 |
| C1CSCCC1(C(=O)O)NC(=O)OCC2C3=CC=CC=C3C4=CC=CC=C24 | AA | acylation | 72 | 82 | 6228.0 | z=-4: 1556, z=-5: 1244.6, z=-6: 1037, z=-7: 888.7, z=-8: 777.5 | 6228.9 |
| C1C[C@H](N(C1)C(=O)OCC2C3=CC=CC=C3C4=CC=CC=C24)CC(=O)O | AA | acylation | 89 | 62 | 6196.0 | z=-4: 1548, z=-5: 1238.2, z=-6: 1031.7, z=-7: 884.1, z=-8: 773.5 | 6198.0 |
| C1C[C@H](N([C@H]1C2=CC=CC=C2)C(=O)OCC3C4=CC=CC=C4C5=CC=CC=C35)C(=O)O | AA | acylation | 85 | 76 | 6258.0 | z=-4: 1563.5, z=-5: 1250.6, z=-6: 1042, z=-7: 893, z=-8: 781.3 | 6258.0 |
| CC(C)(C)OC(=O)N1CCC(CC1)(CC(=O)O)NC(=O)OCC2C3=CC=CC=C3C4=CC=CC=C24 | AA | acylation | 74 | 73 | 6325.1 | z=-4: 1580.3, z=-5: 1264, z=-6: 1053.2, z=-7: 902.6, z=-8: 789.6 | 6325.2 |
| C1=CC=C2C(=C1)C(C3=CC=CC=C32)COC(=O)N[C@@H](CC4=CC=CS4)C(=O)O | AA | acylation | 89 | 70 | 6238.0 | z=-4: 1558.5, z=-5: 1246.6, z=-6: 1038.7, z=-7: 890.1, z=-8: 778.7 | 6238.6 |
| CC(=O)NCSC(C)(C)[C@@H](C(=O)O)NC(=O)OCC1C2=CC=CC=C2C3=CC=CC=C13 | AA | acylation | 76 | 80 | 6287.0 | z=-4: 1570.8, z=-5: 1256.4, z=-6: 1046.8, z=-7: 897.1, z=-8: 784.9 | 6287.1 |
| C1=CC=C2C(=C1)C(C3=CC=CC=C32)COC(=O)N[C@@H](CC4=CSC=N4)C(=O)O | AA | acylation | 89 | 87 | 6239.0 | z=-4: 1558.7, z=-5: 1246.8, z=-6: 1038.8, z=-7: 890.3, z=-8: 778.9 | 6240.3 |
| C[C@H]([C@@H](C(=O)O)NC(=O)OCC1C2=CC=CC=C2C3=CC=CC=C13)OP(=O)(O)OCC4=CC=CC=C4 | AA | acylation | 65 | 16 | 6356.0 | z=-4: 1588, z=-5: 1270.2, z=-6: 1058.3, z=-7: 907, z=-8: 793.5 | 6356.8 |
| C1[C@@H](N(CS1)C(=O)OCC2C3=CC=CC=C3C4=CC=CC=C24)C(=O)O | AA | acylation | 93 | 57 | 6200.0 | z=-4: 1549, z=-5: 1239, z=-6: 1032.3, z=-7: 884.7, z=-8: 774 | 6199.7 |
| C[C@@H]1[C@H](N(C(O1)(C)C)C(=O)[C@H](CC2=CC=CC=C2)NC(=O)OCC3C4=CC=CC=C4C5=CC=CC=C35)C(=O)O | AA | acylation | 81 | 78 | 6373.1 | z=-4: 1592.3, z=-5: 1273.6, z=-6: 1061.2, z=-7: 909.4, z=-8: 795.6 | 6373.2 |
| C1=CC=C(C(=C1)C[C@@H](C(=O)O)NC(=O)OCC2C3=CC=CC=C3C4=CC=CC=C24)C#N | AA | acylation | 90 | 76 | 6257.0 | z=-4: 1563.3, z=-5: 1250.4, z=-6: 1041.8, z=-7: 892.9, z=-8: 781.1 | 6257.7 |
| C1C(N(C2=CC=CC=C21)C(=O)OCC3C4=CC=CC=C4C5=CC=CC=C35)C(=O)O | AA | acylation | 85 | 76 | 6230.0 | z=-4: 1556.5, z=-5: 1245, z=-6: 1037.3, z=-7: 889, z=-8: 777.8 | 6230.3 |
| CN(CC(=O)O)C(=O)OCC1C2=CC=CC=C2C3=CC=CC=C13 | AA | acylation | 91 | 67 | 6156.0 | z=-4: 1538, z=-5: 1230.2, z=-6: 1025, z=-7: 878.4, z=-8: 768.5 | 6155.9 |
| C1CN(CCC1(C2=CC=CC=C2)C(=O)O)C(=O)OCC3C4=CC=CC=C4C5=CC=CC=C35 | AA | acylation | 73 | 75 | 6272.1 | z=-4: 1567, z=-5: 1253.4, z=-6: 1044.3, z=-7: 895, z=-8: 783 | 6273.6 |
| CC1(OC2=C(O1)C=C(C=C2)C[C@@H](C(=O)O)NC(=O)OCC3C4=CC=CC=C4C5=CC=CC=C35)C | AA | acylation | 87 | 90 | 6304.1 | z=-4: 1575, z=-5: 1259.8, z=-6: 1049.7, z=-7: 899.6, z=-8: 787 | 6305.5 |
| C1C2=CC=CC=C2C(N1C(=O)OCC3C4=CC=CC=C4C5=CC=CC=C35)C(=O)O | AA | acylation | 87 | 78 | 6230.0 | z=-4: 1556.5, z=-5: 1245, z=-6: 1037.3, z=-7: 889, z=-8: 777.8 | 6230.5 |
| C1COCCC1(CC(=O)O)NC(=O)OCC2C3=CC=CC=C3C4=CC=CC=C24 | AA | acylation | 73 | 90 | 6226.0 | z=-4: 1555.5, z=-5: 1244.2, z=-6: 1036.7, z=-7: 888.4, z=-8: 777.3 | 6227.1 |
| CC(=O)NCSC[C@@H](C(=O)O)NC(=O)OCC1C2=CC=CC=C2C3=CC=CC=C13 | AA | acylation | 87 | 66 | 6259.0 | z=-4: 1563.8, z=-5: 1250.8, z=-6: 1042.2, z=-7: 893.1, z=-8: 781.4 | 6259.6 |
| CC1=CC=C(C=C1)S(=O)(=O)NC(=NCCC[C@@H](C(=O)O)NC(=O)OCC2C3=CC=CC=C3C4=CC=CC=C24)N | AA | acylation | 87 | 80 | 6395.1 | z=-4: 1597.8, z=-5: 1278, z=-6: 1064.8, z=-7: 912.6, z=-8: 798.4 | 6397.0 |
| CC(C)(C(=O)O)NC(=O)OCC1C2=CC=CC=C2C3=CC=CC=C13 | AA | acylation | 69 | 82 | 6170.0 | z=-4: 1541.5, z=-5: 1233, z=-6: 1027.3, z=-7: 880.4, z=-8: 770.3 | 6170.6 |
| C1CC1C[C@@H](C(=O)O)NC(=O)OCC2C3=CC=CC=C3C4=CC=CC=C24 | AA | acylation | 85 | 73 | 6196.0 | z=-4: 1548, z=-5: 1238.2, z=-6: 1031.7, z=-7: 884.1, z=-8: 773.5 | 6196.2 |
| C1CC(CCC1=O)(C(=O)O)NC(=O)OCC2C3=CC=CC=C3C4=CC=CC=C24 | AA | acylation | 88 | 67 | 6200.0 | z=-4: 1549, z=-5: 1239, z=-6: 1032.3, z=-7: 884.7, z=-8: 774 | 6202.2 |
| CC[C@H](C)[C@@H]([C@H](CC(=O)O)O)NC(=O)OCC1C2=CC=CC=C2C3=CC=CC=C13 | AA | acylation | 84 | 97 | 6242.1 | z=-4: 1559.5, z=-5: 1247.4, z=-6: 1039.3, z=-7: 890.7, z=-8: 779.3 | 6242.4 |
| C1CCC(C1)(C(=O)O)NC(=O)OCC2C3=CC=CC=C3C4=CC=CC=C24 | AA | acylation | 80 | 89 | 6196.0 | z=-4: 1548, z=-5: 1238.2, z=-6: 1031.7, z=-7: 884.1, z=-8: 773.5 | 6197.1 |
| C1[C@H](N(CC2=C1C3=CC=CC=C3N2)C(=O)OCC4C5=CC=CC=C5C6=CC=CC=C46)C(=O)O | AA | acylation | 86 | 57 | 6283.0 | z=-4: 1569.8, z=-5: 1255.6, z=-6: 1046.2, z=-7: 896.6, z=-8: 784.4 | 6283.8 |
| C1=CC=C(C=C1)[C@H]([C@H](C(=O)O)NC(=O)OCC2C3=CC=CC=C3C4=CC=CC=C24)O | AA | acylation | 83 | 83 | 6248.0 | z=-4: 1561, z=-5: 1248.6, z=-6: 1040.3, z=-7: 891.6, z=-8: 780 | 6248.9 |
| C1=CC=C2C(=C1)C(C3=CC=CC=C32)COC(=O)N[C@@H](CC(=O)O)C4=CC=CC=C4C(F)(F)F | AA | acylation | 79 | 58 | 6300.0 | z=-4: 1574, z=-5: 1259, z=-6: 1049, z=-7: 899, z=-8: 786.5 | 6301.5 |
| C1CN(CCC1N2C3=CC=CC=C3N(C2=O)CC(=O)O)C(=O)OCC4C5=CC=CC=C5C6=CC=CC=C46 | AA | acylation | 82 | 79 | 6342.1 | z=-4: 1584.5, z=-5: 1267.4, z=-6: 1056, z=-7: 905, z=-8: 791.8 | 6342.6 |
| C1CC2=C3C(=CC=C2)C[C@H](N3C(=O)[C@H]1NC(=O)OCC4C5=CC=CC=C5C6=CC=CC=C46)C(=O)O | AA | acylation | 90 | 77 | 6313.1 | z=-4: 1577.3, z=-5: 1261.6, z=-6: 1051.2, z=-7: 900.9, z=-8: 788.1 | 6313.4 |
| C1C[C@H]2C[C@@H]1[C@@H]([C@@H]2NC(=O)OCC3C4=CC=CC=C4C5=CC=CC=C35)C(=O)O | AA | acylation | 83 | 87 | 6222.0 | z=-4: 1554.5, z=-5: 1243.4, z=-6: 1036, z=-7: 887.9, z=-8: 776.8 | 6222.3 |
| C1C2=CC=CC=C2CC1(C(=O)O)NC(=O)OCC3C4=CC=CC=C4C5=CC=CC=C35 | AA | acylation | 77 | 63 | 6244.0 | z=-4: 1560, z=-5: 1247.8, z=-6: 1039.7, z=-7: 891, z=-8: 779.5 | 6243.9 |
| C1=CC=C2C(=C1)C(C3=CC=CC=C32)COC(=O)N[C@@H](CC4=CNC5=CC=CC=C54)C(=O)O | AA | acylation | 87 | 76 | 6271.0 | z=-4: 1566.8, z=-5: 1253.2, z=-6: 1044.2, z=-7: 894.9, z=-8: 782.9 | 6272.0 |
| C1=CC=C2C(=C1)C(C3=CC=CC=C32)COC(=O)N[C@@H](CC4=CC=C(C=C4)O)C(=O)O | AA | acylation | 90 | 73 | 6248.0 | z=-4: 1561, z=-5: 1248.6, z=-6: 1040.3, z=-7: 891.6, z=-8: 780 | 6248.3 |

**Table S7.** Chemical modification of immobilized nucleic acid. 500 pmol of a model $NH_2$-modified nucleic acid was immobilized on 1-µm magnetic DEAE WAX resin and acylated with 4-formylbenzoic acid followed by reductive amination. After elution from the resin, samples were analyzed by HPLC (260 nm). Conversion (96%) was calculated by comparison of the expected product peak area to the starting material. Product identification was confirmed by MS. Purity of the product (100%) was calculated as a percentage of all peaks excluding any unconverted starting material.

| SMILES | Monomer Class | Reaction | Conversion (%) | Purity (%) | Theoretical MW | Expected m/z | Calculated MW |
|---|---|---|---|---|---|---|---|
| CC(=O)NCCN | AM | reductive_amination | 94 | 85 | 6081.1 | z=-4: 1519.3, z=-5: 1215.2, z=-6: 1012.5, z=-7: 867.7, z=-8: 759.1 | 6081.4 |
| NCC1CCCCC1 | AM | reductive_amination | 88 | 95 | 6092.1 | z=-4: 1522, z=-5: 1217.4, z=-6: 1014.4, z=-7: 869.3, z=-8: 760.5 | 6092.9 |
| NC[C@H]1N(C)CCC1 | AM | reductive_amination | 100 | 86 | 6093.1 | z=-4: 1522.3, z=-5: 1217.6, z=-6: 1014.5, z=-7: 869.4, z=-8: 760.6 | 6092.9 |
| NCCN1CCOCC1 | AM | reductive_amination | 100 | 96 | 6109.1 | z=-4: 1526.3, z=-5: 1220.8, z=-6: 1017.2, z=-7: 871.7, z=-8: 762.6 | 6108.2 |
| NCCC1=CC=CO1 | AM | reductive_amination | 100 | 85 | 6090.1 | z=-4: 1521.5, z=-5: 1217, z=-6: 1014, z=-7: 869, z=-8: 760.3 | 6091.1 |
| NCCN1C=CN=C1 | AM | reductive_amination | 87 | 93 | 6090.1 | z=-4: 1521.5, z=-5: 1217, z=-6: 1014, z=-7: 869, z=-8: 760.3 | 6090.2 |
| NCC1=CC=CC=C1 | AM | reductive_amination | 94 | 93 | 6086.1 | z=-4: 1520.5, z=-5: 1216.2, z=-6: 1013.3, z=-7: 868.4, z=-8: 759.8 | 6086.1 |
| OC1=CC=C(CN)C=C1 | AM | reductive_amination | 100 | 94 | 6102.1 | z=-4: 1524.5, z=-5: 1219.4, z=-6: 1016, z=-7: 870.7, z=-8: 761.8 | 6102.3 |
| N#CC1=CC=C(CN)C=C1 | AM | reductive_amination | 93 | 90 | 6111.1 | z=-4: 1526.8, z=-5: 1221.2, z=-6: 1017.5, z=-7: 872, z=-8: 762.9 | 6111.6 |
| NCCOC1=CC=CC=C1 | AM | reductive_amination | 98 | 85 | 6116.1 | z=-4: 1528, z=-5: 1222.2, z=-6: 1018.4, z=-7: 872.7, z=-8: 763.5 | 6116.4 |
| NC[C@H]1CN(C2=CC=C(N3CCOCC3=O)C=C2)C(O1)=O | AM | reductive_amination | 100 | 100 | 6270.2 | z=-4: 1566.6, z=-5: 1253, z=-6: 1044, z=-7: 894.7, z=-8: 782.8 | 6270.1 |

#### Supplemental Figures

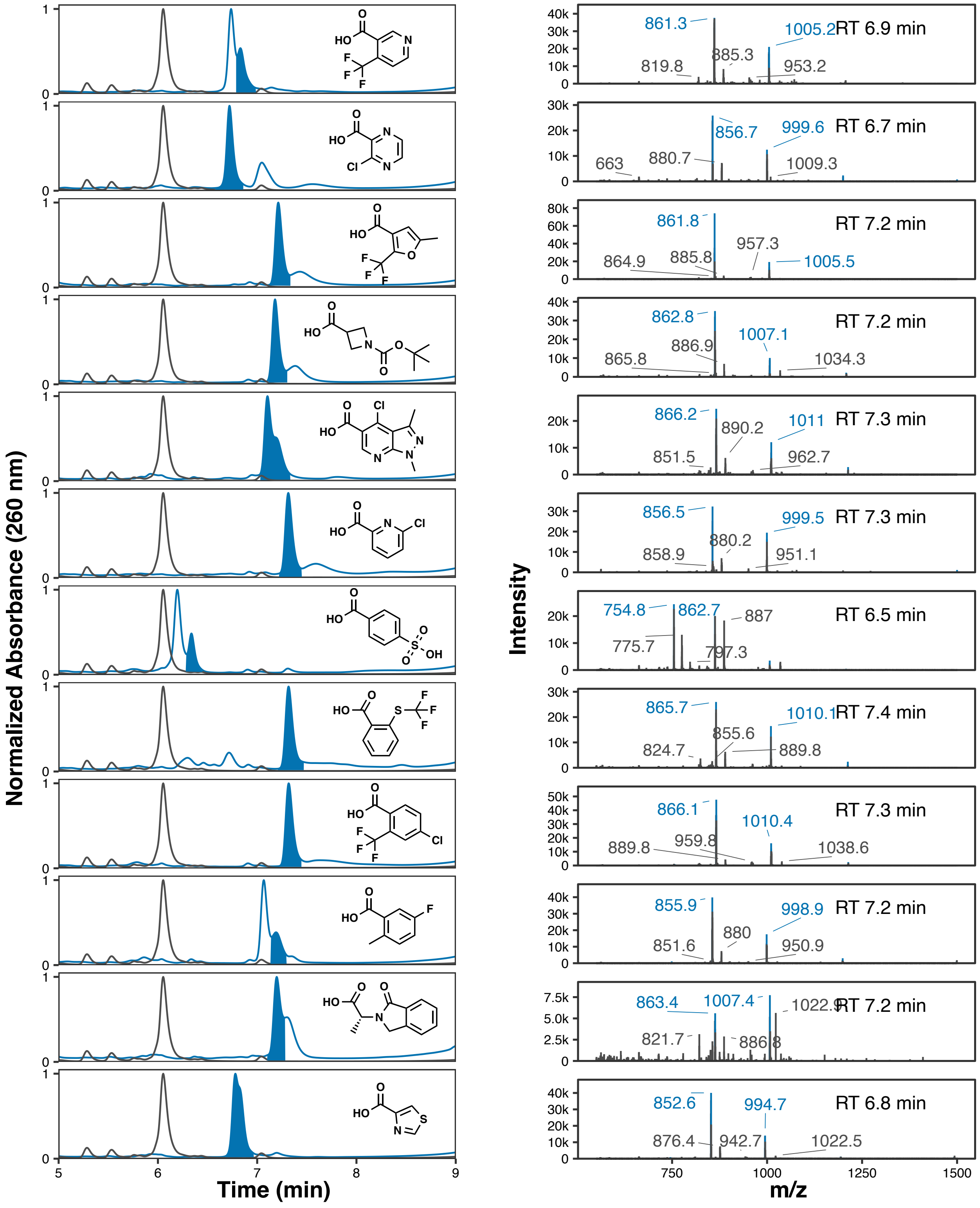

**Figure S1.** LC-MS analysis of WAX-ON acylation of DNA-$NH_2$ with various CA BBs. A model 5′-$NH_2$-modified DNA oligonucleotide was immobilized on 1-µm magnetic DEAE WAX resin and acylated with CA BBs used for DEL construction. After elution from the resin, samples were analyzed by LC-MS and the area of the modified peak (blue) compared to the area of any unmodified oligo (black trace). Product identification was confirmed by MS; expected ions of a calculated charge state envelope are highlighted (blue).

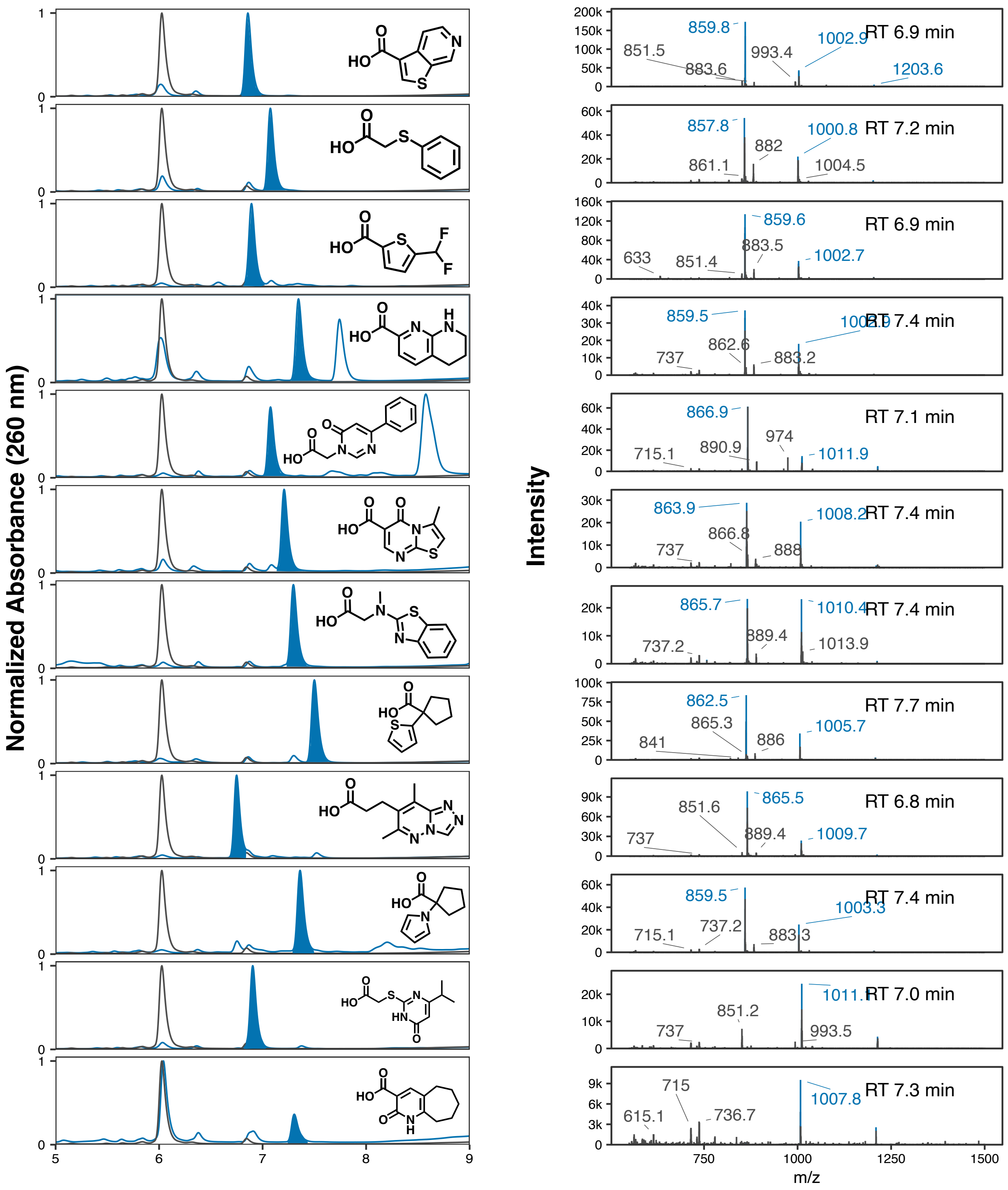

**Figure S1.** LC-MS analysis of WAX-ON acylation of DNA-$NH_2$ with various CA BBs. A model 5′-$NH_2$-modified DNA oligonucleotide was immobilized on 1-µm magnetic DEAE WAX resin and acylated with CA BBs used for DEL construction. After elution from the resin, samples were analyzed by LC-MS and the area of the modified peak (blue) compared to the area of any unmodified oligo (black trace). Product identification was confirmed by MS; expected ions of a calculated charge state envelope are highlighted (blue).

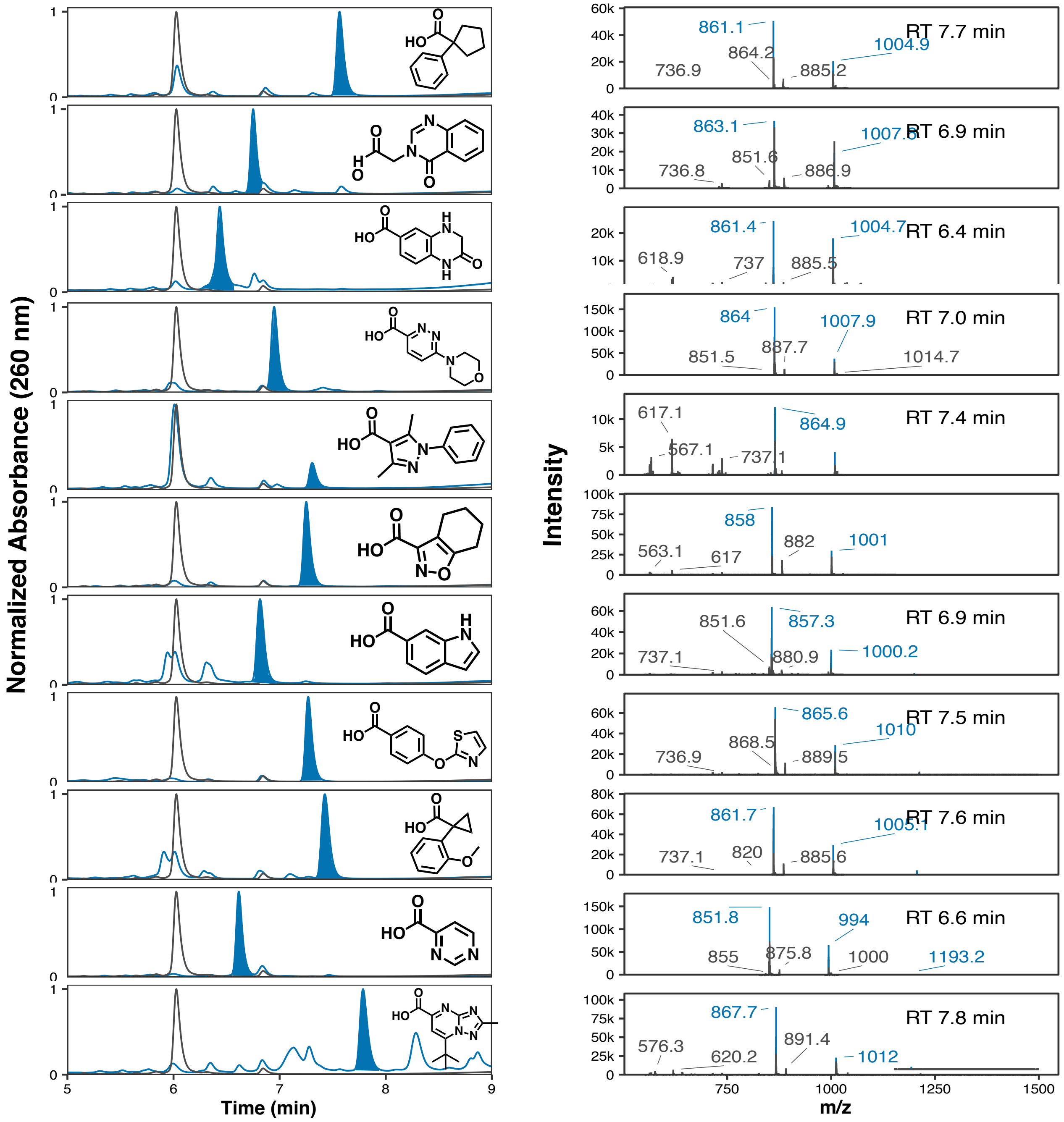


**Figure S1.** LC-MS analysis of WAX-ON acylation of DNA-$NH_2$ with various CA BBs. A model 5′-$NH_2$-modified DNA oligonucleotide was immobilized on 1-μm magnetic DEAE WAX resin and acylated with CA BBs used for DEL construction. After elution from the resin, samples were analyzed by LC-MS and the area of the modified peak (blue) compared to the area of any unmodified oligo (black trace). Product identification was confirmed by MS; expected ions of a calculated charge state envelope are highlighted (blue).

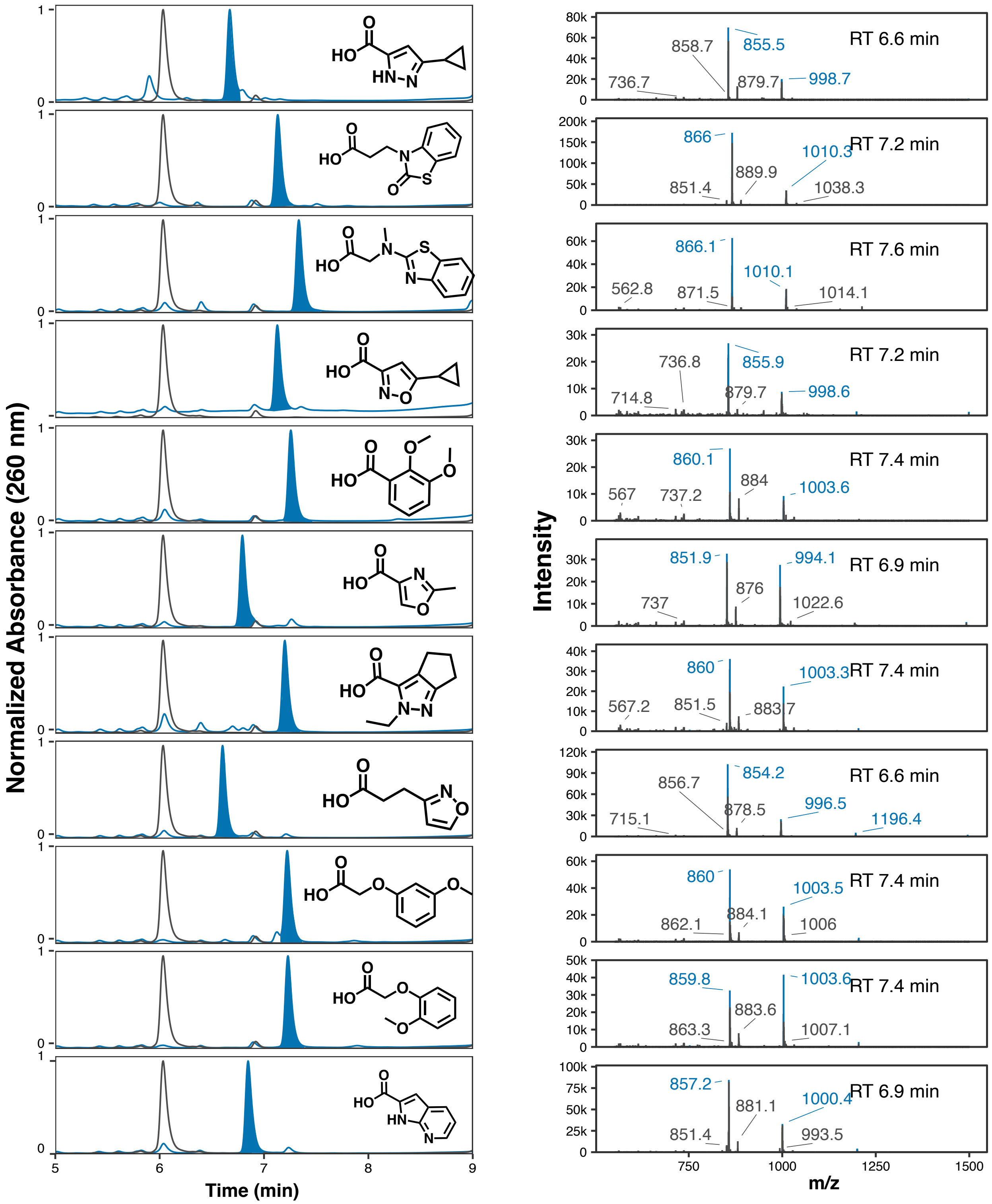


**Figure S1.** LC-MS analysis of WAX-ON acylation of DNA-$NH_2$ with various CA BBs. A model 5′-$NH_2$-modified DNA oligonucleotide was immobilized on 1-µm magnetic DEAE WAX resin and acylated with CA BBs used for DEL construction. After elution from the resin, samples were analyzed by LC-MS and the area of the modified peak (blue) compared to the area of any unmodified oligo (black trace). Product identification was confirmed by MS; expected ions of a calculated charge state envelope are highlighted (blue).

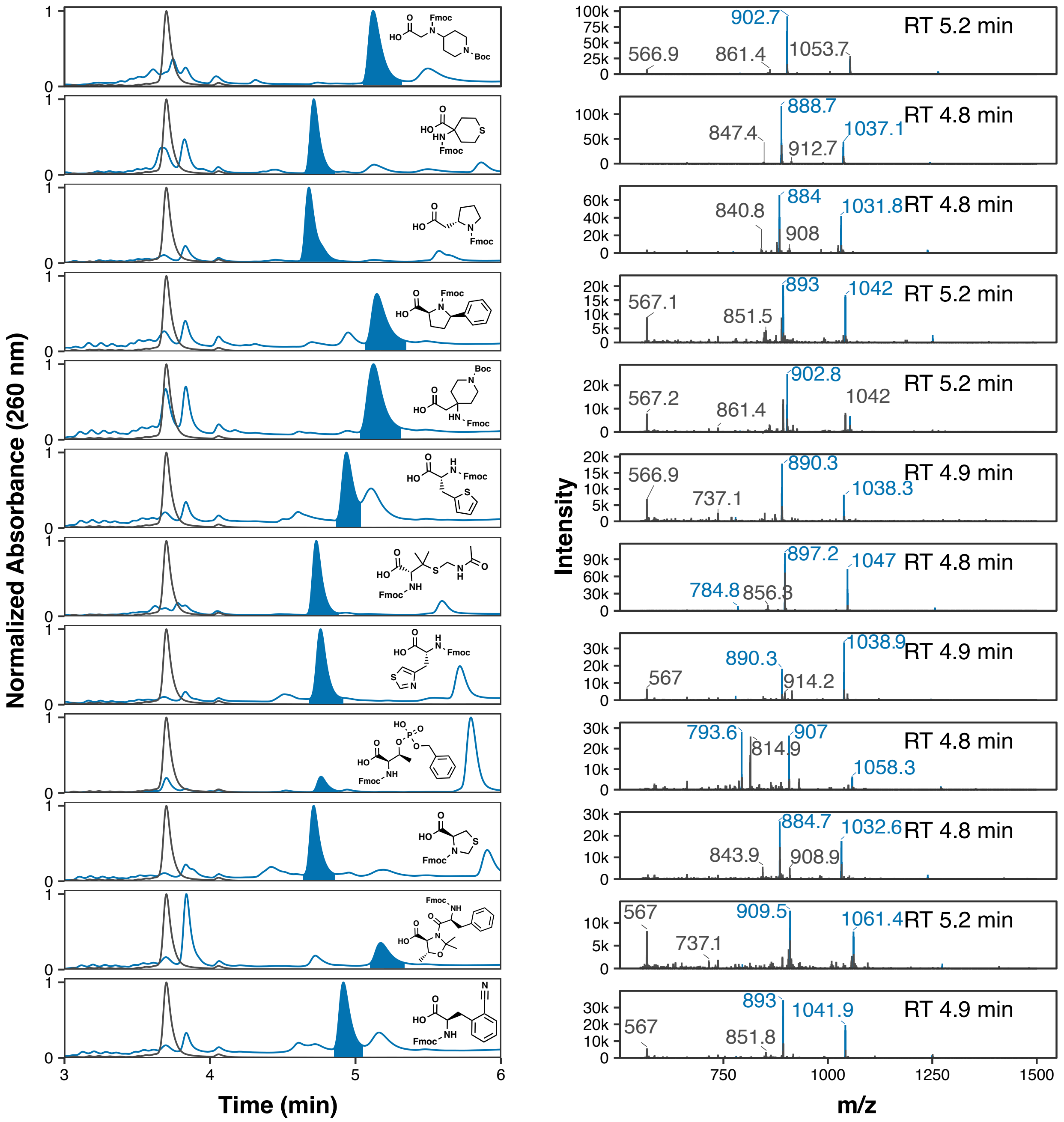


**Figure S2.** LC-MS analysis of WAX-ON acylation of DNA-$NH_2$ with various AA BBs. A model 5′-$NH_2$-modified DNA oligonucleotide was immobilized on 1-μm magnetic DEAE WAX resin and acylated with AA BBs used for DEL construction. After elution from the resin, samples were analyzed by HPLC and the area of the modified peak (blue) compared to the area of any unmodified oligo (black trace). Product identification was confirmed by MS; expected ions of a calculated charge state envelope are highlighted (blue).

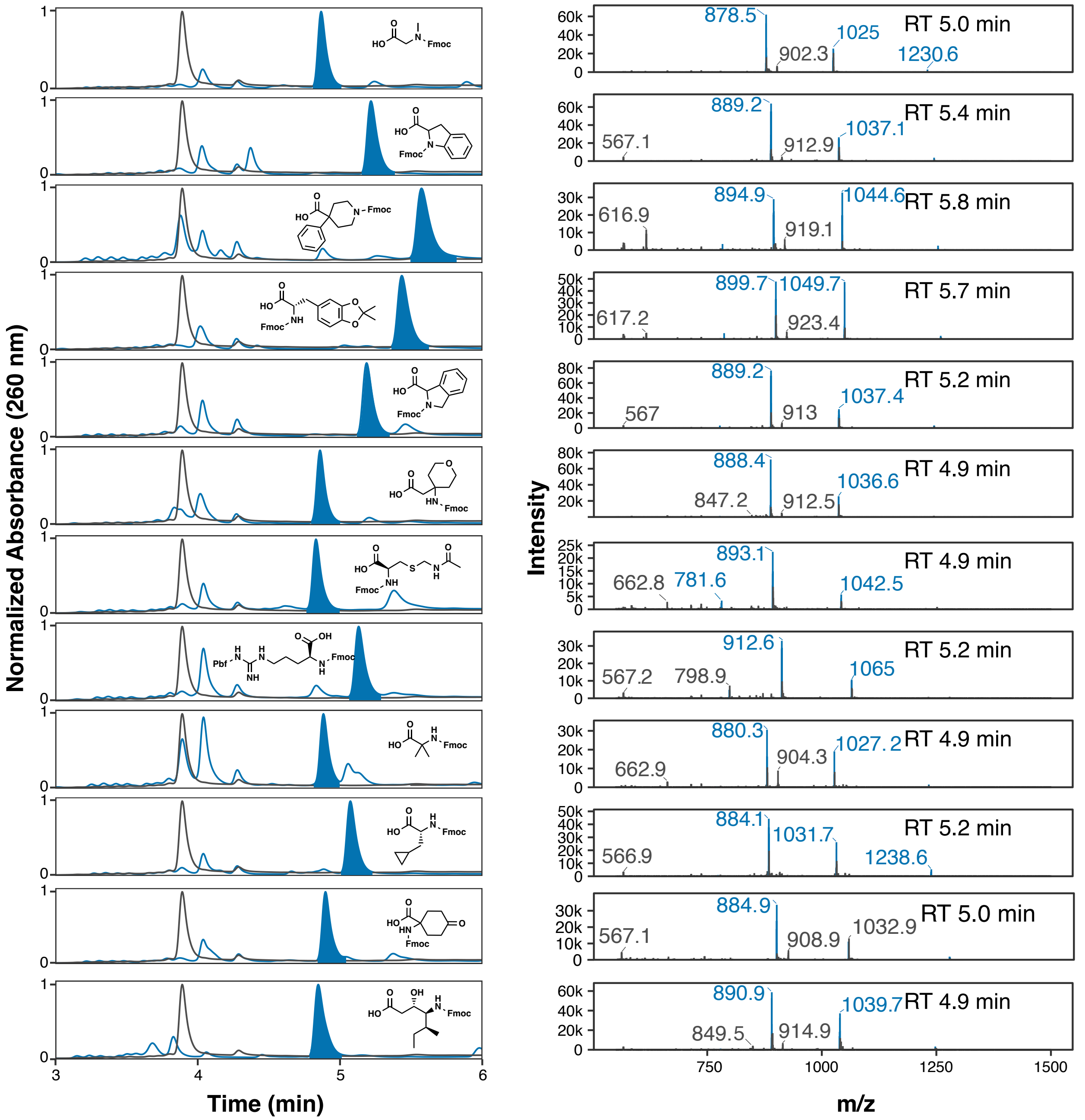


**Figure S2.** LC-MS analysis of WAX-ON acylation of DNA-$NH_2$ with various AA BBs. A model 5′-$NH_2$-modified DNA oligonucleotide was immobilized on 1-µm magnetic DEAE WAX resin and acylated with AA BBs used for DEL construction. After elution from the resin, samples were analyzed by HPLC and the area of the modified peak (blue) compared to the area of any unmodified oligo (black trace). Product identification was confirmed by MS; expected ions of a calculated charge state envelope are highlighted (blue).

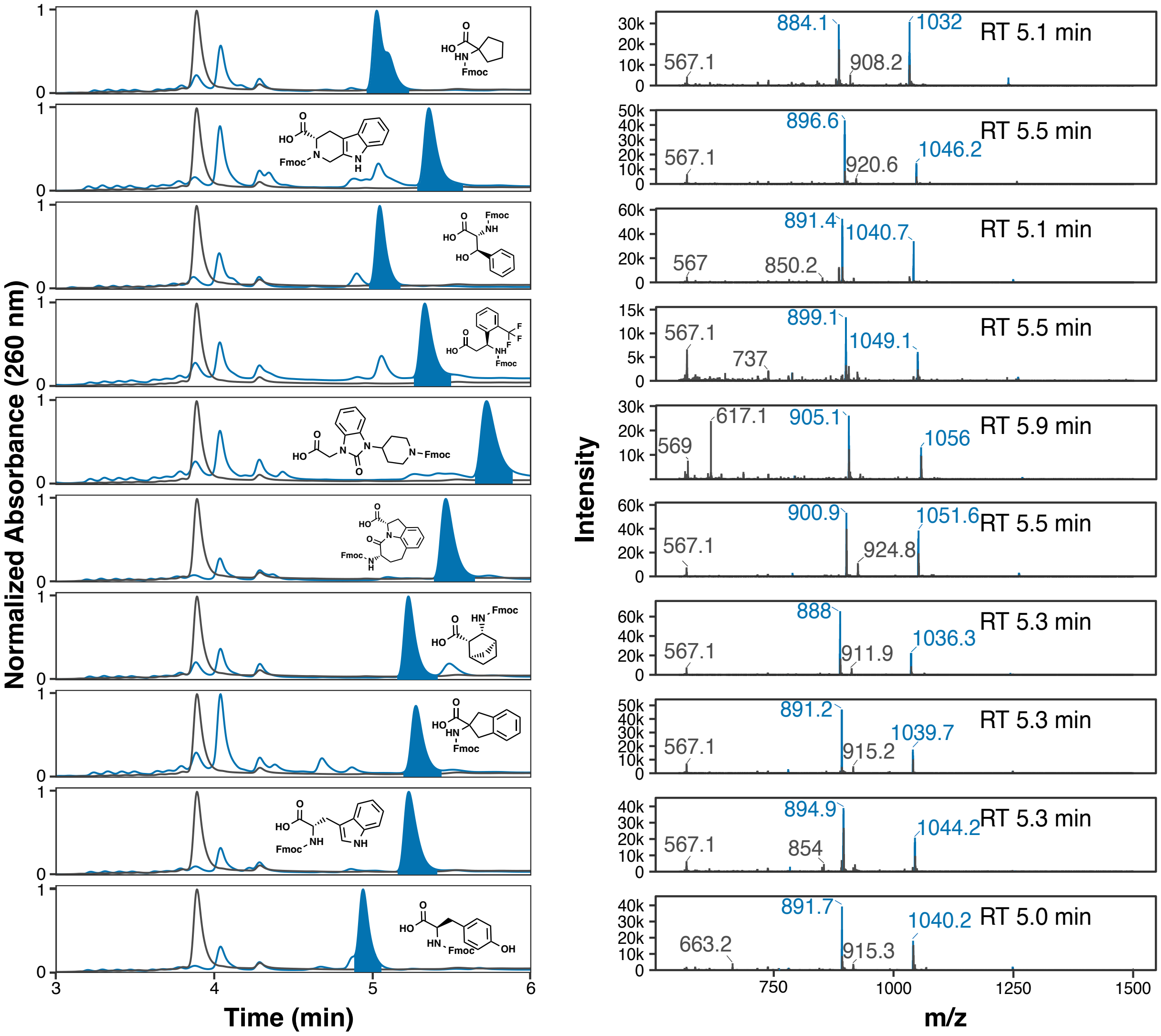


**Figure S2.** LC-MS analysis of WAX-ON acylation of DNA-$NH_2$ with various AA BBs. A model 5′-$NH_2$-modified DNA oligonucleotide was immobilized on 1-µm magnetic DEAE WAX resin and acylated with AA BBs used for DEL construction. After elution from the resin, samples were analyzed by HPLC and the area of the modified peak (blue) compared to the area of any unmodified oligo (black trace). Product identification was confirmed by MS; expected ions of a calculated charge state envelope are highlighted (blue).

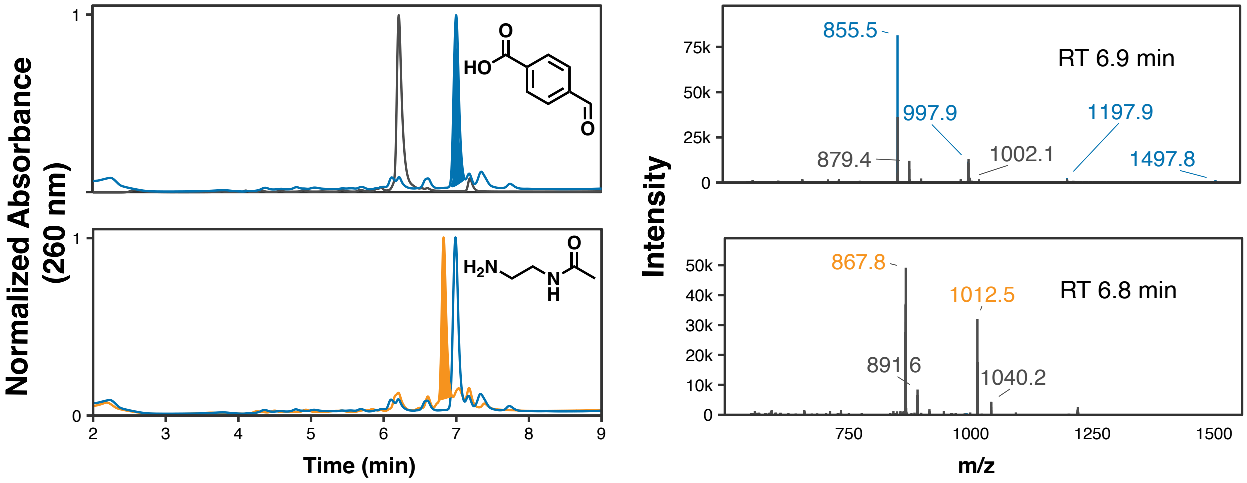


**Figure S3.** A model 5′-$NH_2$-modified DNA oligonucleotide was immobilized on 1-µm magnetic DEAE WAX resin and acylated with 4-formylbenzoic acid (blue). The aldehyde was subsequently coulped via reductive amination with an Am BB (N-acetylethylenediamine, orange) while still immobilized on resin. After elution from the resin, samples were analyzed by HPLC and the area of the modified peak (blue) compared to the area of any unmodified oligo (black trace). Product identification was confirmed by MS; expected ions of a calculated charge state envelope are highlighted.

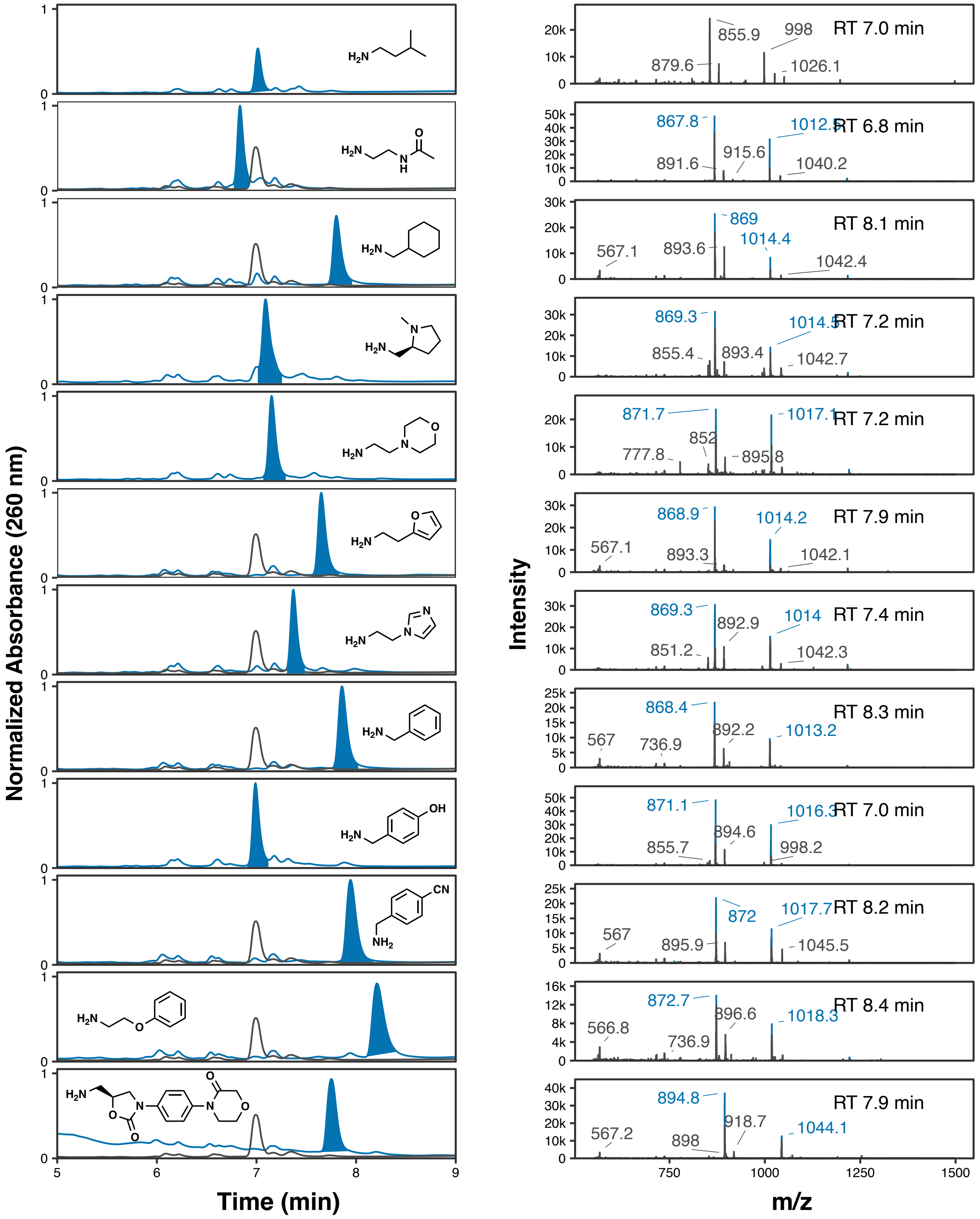


**Figure S4.** Reductive amination of nucleic acid immobilized on magnetic DEAE WAX resin. A model 5′-$NH_2$-modified DNA oligonucleotide was immobilized on 1-µm magnetic DEAE WAX resin, acylated with 4-formylbenzoic acid, and then reacted with various Am BBs via reductive amination. After elution from the resin, samples were analyzed by HPLC and the area of the modified peak (blue) compared to the area of the acylated oligo (black trace). Product identification was confirmed by MS; expected ions of a calculated charge state envelope are highlighted (blue).

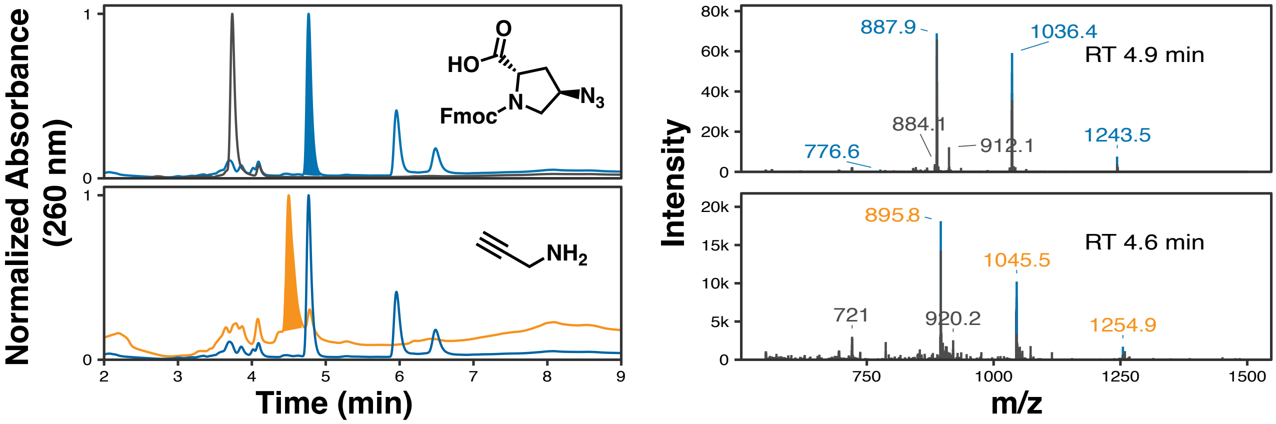


**Figure S5.** CuAAC of nucleic acid immobilized on magnetic DEAE WAX resin. A model 5′-$NH_2$-modified DNA oligonucleotide (black) was immobilized on 1-µm magnetic DEAE WAX resin and acylated with Fmoc-$N_3$-Pro-OH (blue). The immobilized material was subsequently modified with an alkyne (propargylamine, orange) via CuAAC reaction. Product identification was confirmed by MS; expected ions of the calculated charge state envelopes are highlighted.
